# Non-viral gene editing *in utero* with lipid nanoparticles complexed to mRNA

**DOI:** 10.1101/2022.10.14.512310

**Authors:** Kewa Gao, Jie Li, Hengyue Song, Hesong Han, Yongheng Wang, Boyan Yin, Diana L. Farmer, Niren Murthy, Aijun Wang

## Abstract

Nanoparticle-based drug delivery systems have the potential to revolutionize medicine but their low vascular permeability and rapid clearance by phagocytic cells have limited their medical impact. Nanoparticles delivered at the *in utero* stage have the potential to overcome these key limitations, due to the high rate of angiogenesis and cell division in fetal tissue, and the under-developed immune system. However, very little is known about nanoparticle drug delivery at the fetal stage of development. In this report, using Ai9 CRE reporter mice, we demonstrate that lipid nanoparticle (LNP) mRNA complexes can deliver mRNA for gene editing enzymes *in utero* after an intrahepatic injection, and can access and edit major organs, such as the heart, the liver, kidneys, lungs and the gastrointestinal tract with remarkable efficiency and low toxicity. In addition, we show here that Cas9 mRNA and sgRNA complexed to LNPs were able to edit the fetal organs *in utero* after an intrahepatic injection. These experiments demonstrate the possibility of non-viral delivery of gene editing enzymes *in utero* and nanoparticle drug delivery has great potential for delivering macromolecules to organs outside of the liver *in utero*, which provides a promising strategy for treating a wide variety of devastating genetic diseases before birth.

## Introduction

Gene editing therapeutics have the potential to revolutionize the treatment of genetic diseases. However, delivery challenges have limited their clinical translation[1, 2]. Delivering gene editing enzymes *in vivo* is challenging because delivery vehicles for these enzymes, such as lipid nanoparticles (LNPs), viruses or polymer nanoparticles are all greater than 15 nm and cannot extravasate from the blood into the tissues[3–5]. Consequently, intravenous injections of gene editing delivery vehicles generally result in only editing of the liver and other phagocytic organs. However, the vast majority of genetic diseases require treatments that are based upon transfecting non-liver organs such as the kidneys, heart, lungs, GI tract and brain, which require extravasation from the blood and therefore cannot be transfected via delivery vectors greater than 15 nm [3]. The extravasation of gene editing enzymes from the blood into tissues is a central problem in gene editing and will require fundamentally new drug delivery approaches.

Gene editing at the *in utero* stage has the potential to address several of the delivery challenges associated with editing non-phagocytic tissues and has great therapeutic potential. For example, a human fetus in the second trimester is about 1% the weight of a 1-year-old child [6], and this is a gestational age at which systemic delivery of macromolecules is technically feasible. In addition, the fetal immune system is tolerogenic and may not generate an immune response against foreign gene editing enzymes and immunostimulatory nanoparticles, which is a frequent problem in adults [6, 7]. Additionally, stem and progenitor cells are prevalent in the developing fetus and often actively proliferate, which may facilitate more efficient gene editing [8, 9]. Moreover, the barriers associated with blood vessel permeability may not be present and phagocytosis is premature at the developing fetal stage, therefore systemically applied nanoparticle-based reagents have the potential to transfect and edit multiple organs such as the liver, heart, skin, GI tract and brain at the *in utero* stage [7, 10–12]. In addition, *in utero* injections can be performed safely, at low cost without the need for sophisticated infrastructure, and can be done in all parts of the world including the economically under-developed regions [8]. Finally, *in utero* editing does not cause germ-line editing and therefore avoids the ethical dilemmas associated with germline editing [13–15]. Therefore, *in utero* gene editing has the potential to transform the treatment of genetic diseases. However, before *in utero* gene editing is possible, safe and effective, methods for delivering gene editing enzymes *in utero* must be developed.

Several strategies for performing *in utero* gene editing are currently being explored. Adenoviral or adeno-associated virus delivery of gene editing enzymes *in utero* has been accomplished with good editing efficiency[7, 11], but viral-based methods are problematic because they lead to sustained expression of gene editing enzymes, which could cause high levels of off-target DNA damage[16–18]. Non-viral gene editing has also been explored via the delivery of peptide nucleic acids (PNAs) in PLGA nanoparticles [8]. However, it is unclear if this methodology extends towards gene editing enzymes. These pioneering studies demonstrated that gene editing *in utero* was possible and could be accomplished with low toxicity.

mRNA/LNP complexes are a potentially powerful methodology for transfecting gene editing enzymes *in vivo* because of the highly modular nature of mRNA[19] and the low toxicity of LNPs[20] in comparison to viral-based delivery methods. The feasibility of delivering mRNA *in utero* has recently been demonstrated by Riley et al, who demonstrated that an intravascular delivery of LNP mRNA complexes could transfect cells in the liver with an efficiency of approximately 1%[9]. These pioneering studies have demonstrated that LNPs containing GFP mRNA are well tolerated *in utero* after an intravascular delivery. However, it is still unclear if gene editing enzymes can be delivered *in utero* via mRNA-based delivery methods, and what cell types will be transfected *in utero* with mRNA/LNP complexes. In addition, it is likely that a transfection efficiency > 1% will be needed for clinical applications.

In this report, using Ai9 CRE reporter mice, we demonstrate that mRNA/LNP complexes can deliver mRNA for gene editing enzymes *in utero*, and can edit major organs, such as the liver, the heart, kidneys, lungs, GI tract and brain with remarkable efficiency. We show here that CRE mRNA complexed to LNPs were able to edit Ai9 mice *in utero* after an intrahepatic injection, and edited 0.64-12.4% of the cells in the heart, lungs, liver, kidneys, GI tract and brain, with low toxicity. In addition, the mRNA/LNP complex showed a high affinity for muscle tissue, and we found that a large number of muscle stem cells were transfected in the heart, diaphragm and skeletal muscle. At 4 weeks after birth, we demonstrate that 50.99 ± 5.05%, 36.62 ± 3.42% and 23.7 ± 3.21% of muscle fiber in the diaphragm, heart and skeletal muscle were transfected respectively. In addition, Cas9 mRNA and sgRNA complexed to LNPs, were able to also edit Ai9 mice *in utero* after an intrahepatic injection, and edited 0.2-0.8% of cells in the liver, heart, GI, and kidneys. These experiments are demonstration of non-viral delivery of gene editing enzymes *in utero* and demonstrate that this therapeutic approach has tremendous therapeutic potential.

## Results and Discussion

### Scientific rationale for intrahepatic injection of gene editing enzymes *in utero*

Delivering nanoparticles to organs outside of the liver is a major challenge in adults due to the lower permeability of the vasculature, the large diffusion distance and rapid phagocytic clearance of nanoparticles. The *in utero* environment is dramatically different than in adults, and may offer unique opportunities for nanoparticle drug delivery. In particular, endothelial cells are undergoing significant levels of angiogenesis in all organs *in utero,* and there is consequently great potential for large increases in permeability. Angiogenesis rarely occurs in adults, except in tumors, and nanoparticle drug delivery to tumors has been particularly successful. Similarly, the *in utero* environment may have lower numbers of mature macrophages, and consequently, the circulation half-life of nanoparticles may also be significantly extended.

However, despite its promise, very little is known about *in utero* gene editing thus far. mRNA/LNP complexes have been investigated via intravascular delivery in a murine model with moderate success [9]. Developing effective transfection and delivery approaches to achieve clinically meaningful gene editing are therefore greatly needed. To obtain direct vascular access, intrahepatic *in utero* injection has been well established and can be easily translated to intravenous umbilical cord injection under ultrasound guidance for human patients in clinical settings [21, 22]. Intrahepatic delivery of mRNA/LNP complexes has potential for transfecting the developing fetus, due to the high permeability of the developing organs, and the potential for diffusion to multiple organs simultaneously via a combination of fickian diffusion and transport via vasculature. Compared to reported intravascular delivery in mice, intrahepatic delivery can be carried out at an earlier gestational stage and this may lead to substantial differences in the overall transfection efficacy as well as in the types of cells transfected [21–23].

**Figure 1.**
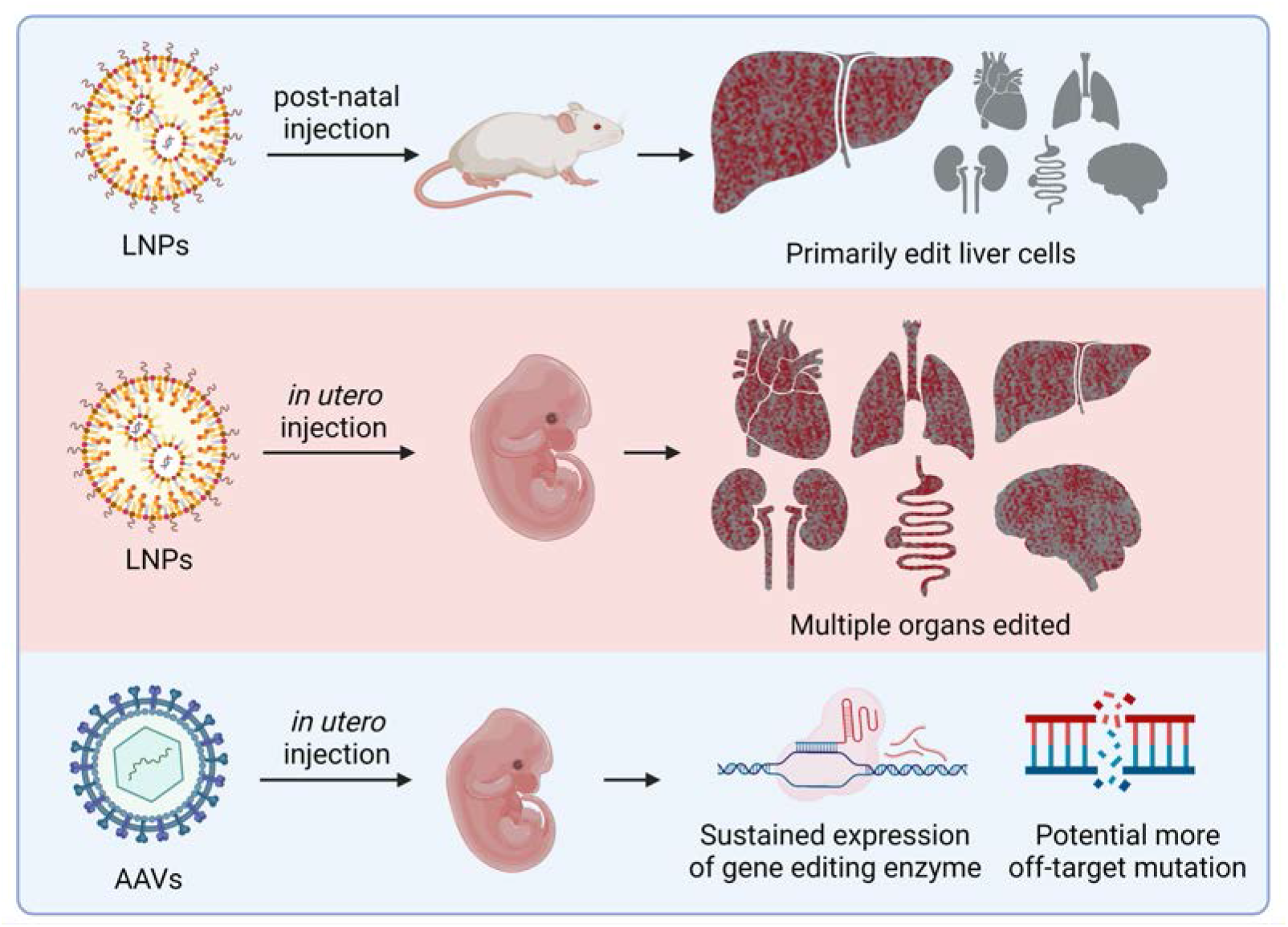
LNPs can deliver mRNA coding for gene editing enzymes *in utero* and efficiently transfect the heart, lung, kidney, liver and GI tract. In this report we demonstrate that lipid nanoparticle (LNP) mRNA complexes can deliver mRNA for gene editing enzymes *in utero* after an intrahepatic injection, and can access and edit major organs, such as the heart, the liver, kidneys, lungs and the gastrointestinal tract with remarkable efficiency. In contrast, in adults the liver is the predominant organ edited when LNPs containing gene editing enzymes are administered. LNP/mRNA complexes coding for gene editing enzymes have potential benefits over AAV based delivery strategies because of their transient nature, and will have lower off-target DNA damage in comparison to AAV. These experiments demonstrate that gene editing enzymes can be delivered *in utero* via non-viral delivery methods and provide a strategy for treating a wide variety of devastating genetic diseases.

### Characterization of LNPs and *in vitro* gene editing efficiency in Ai9 NIH 3T3 cells

LNPs have been widely used for delivery of siRNA and mRNA to treat adult diseases[24, 25] and D-Lin containing LNPs in particular, has been approved by the FDA for treatment of hereditary TTR amyloidosis[25]. D-Lin containing LNPs have been delivered via the intravenous route, *in utero*, with moderate success, however their ability to deliver mRNA via the intrahepatic route has never been investigated. Here we developed 4 different LNPs using D-Lin as ionizable lipid components to explore their potential to deliver RNAs *in utero*. D-Lin containing LNPs were formed using the ethanol dilution method [26] by rapid mixing of lipid mixtures containing ionizable lipid (D-Lin-MC3-DMA), DOPE, cholesterol and PEG-lipids in the ethanol phase with mRNA in the aqueous phase. D-Lin enables mRNA encapsulation, cellular uptake and endosomal escape of the encapsulated mRNA into the cytosol [27], while DOPE and cholesterol provide stability of LNPs and may assist in endosomal escape [28, 29]. The PEG-lipids enhance overall LNP stability and extend circulation [30]. Four different cationic lipid formulations and lipofectamine were screened (see table 1 for details of formulations). LNPs were characterized by size and mRNA encapsulation efficiency (Table S1). The hydrodynamic diameter, as assessed by dynamic light scattering (DLS) for all LNP formulations ranged from 112.5 to 144.0 nm with PDI values less than 0.15, indicating monodisperse LNPs. Each LNP formulation was evaluated for its ability to encapsulate mRNA by RiboGreen assays [31], and all encapsulation efficiencies were high, ranging from 74.6 to 82.9%. The LNPs containing CRE mRNA exhibited high gene editing efficiency in Ai9 NIH 3T3 cells via flow cytometry measurement of edited td-Tomato positive cells. LNPs treated cells showed high gene editing efficiency with td-Tomato positive rate ranging from 53.2%-61.8%, which is at least 66% increase compared with mRNA/lipofectamine 2000 complex (Figure S1). The good mRNA encapsulation efficiency, excellent size distribution and *in vitro* gene editing efficiency of these LNPs indicates that they were good candidates for *in utero* gene editing.

**Table 1.** Detailed formulation components of LNPs.

|  | LNP1 | LNP2 | LNP3 | LNP4 |
| --- | --- | --- | --- | --- |
| D-Lin-MC3-DMA | 36.8% | 45.4% | 38.3% | 35% |
| DOPE | 23.8% | 15.7% | 11.0% | 16% |
| Cholesterol | 38.2% | 37.7% | 49.5% | 46.5% |
| DMG-PEG2000 | 1.2% | 1.2% | 1.2% | 2.5% |

### LNPs can distribute to various internal organs after *in utero* intrahepatic injection

A key challenge with using nanoparticles for drug delivery is their rapid clearance by the liver and their low vascular permeability. However, it is unknown if the *in utero* environment will be similarly challenging for nanoparticle drug delivery. We therefore performed experiments to determine the biodistribution of LNPs injected intrahepatically at the gestational age of E15.5 and compared their biodistribution with LNPs injected in adult mice. For these experiments, we used LNPs that contained Cy7 labelled PEG-Lipid and CRE mRNA. Blue food dye was added to make the injection position visible through the maternal uterus wall and fetal abdominal wall. The blue reagent gathered in the fetal liver right after injection, and then spread throughout the whole fetus with the blood circulation (see surgery procedure in Figure 2B), demonstrating that our injection was successfully delivered into the fetal liver. Saline was injected into fetuses via the same route as LNPs to serve as a negative control.

**Figure 2.**
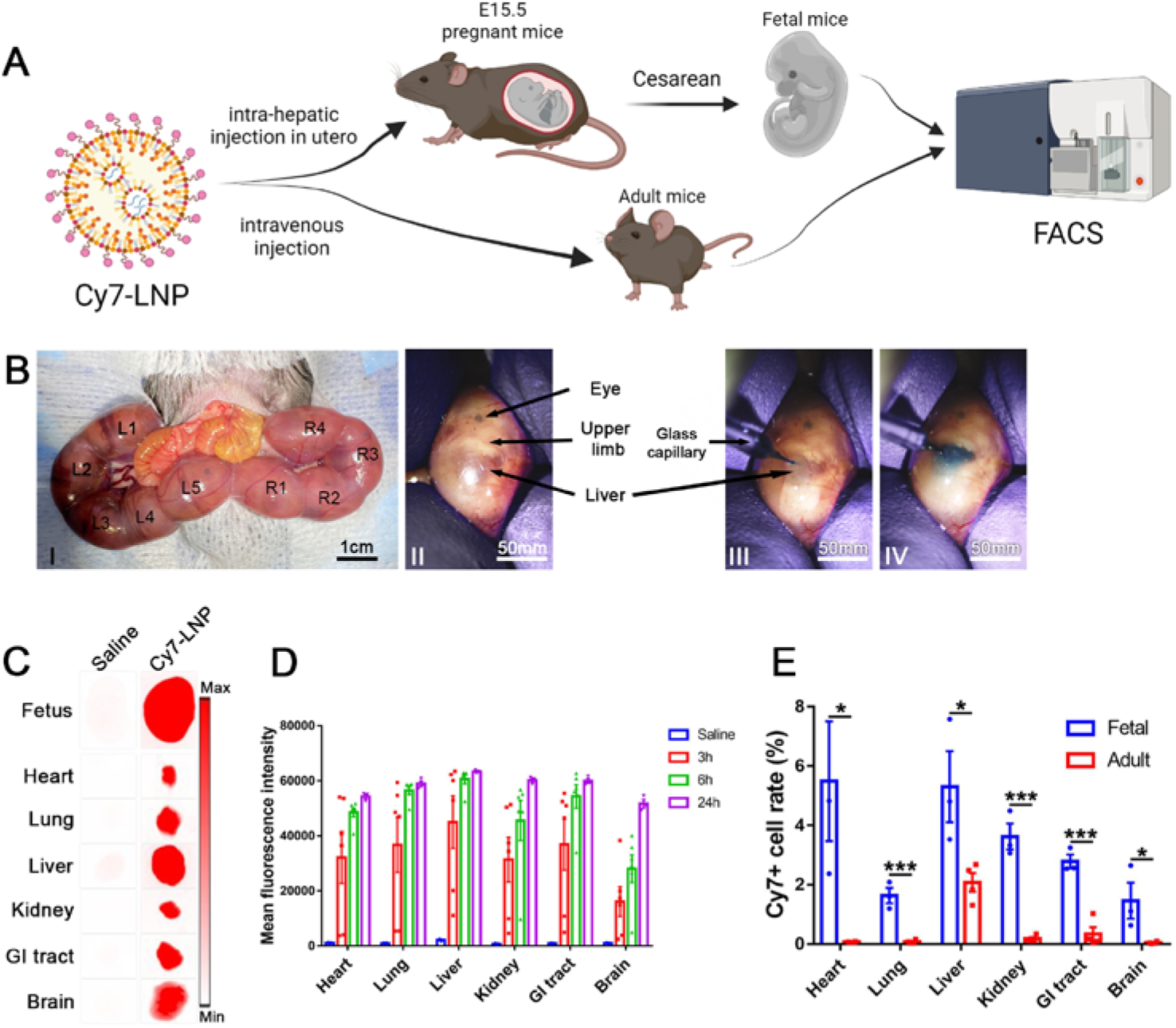
LNPs injected *in utero* are permeable to the vasculature and can distribute to various internal organs. (A) Schematic depicting biodistribution experiments of Cy7 labeled LNP1. (B) Intraoperative photography of the *in utero* intrahepatic injection procedure. (I). The uterus of the pregnant mouse was exposed. (II). Identify the location of the fetal liver. (III). Puncture the fetal liver with a glass capillary. (IV). LNP1 labeled with food dye was injected into the fetal liver. (C) Cy7 labeled LNP1 was imaged in the whole E15.5 mouse fetus and internal organs after *in utero* injection demonstrate that LNP1 can distribute into multiple organs at 24h post *in utero* intrahepatic injection. (D) Quantitative analysis of the fluorescence intensity of Cy7 labeled LNP1. Cy7 signals can already be detected in various organs 3 hours post injection and gradually increase as time increases (Data are represented as mean ± SEM, n = 6). (E) The Cy7 uptake rates post *in utero* injection were measured by flow cytometry and compared to adulthood injection. LNP1 injected during adulthood is mainly captured by cells in the liver. In contrast, LNP1 injected during the fetal period can reach various internal organs after 24 hours post injection. The uptake rates of LNP1 injected during the fetal period was significantly higher than that of LNP1 injected during the adult period. (Data are represented as mean ± SEM, n = 3, *p < 0.05, ***p < 0.0001).

Figures 2C and Figure 2D demonstrate that LNPs are distributed into several internal organs after an *in utero* intrahepatic injection, 3 hours post injection, and gradually increased over time. The Cy7 signal was evenly distributed in the heart, lung, liver, kidney, GI tract and brain, suggesting that the LNPs are accessing the blood after an intrahepatic injection and persisting in the blood for several hours. The proportion of cells that took up Cy7 labeled LNPs in each organ was quantified by flow cytometry. The heart and liver revealed the highest uptake rate of injected LNPs at 24 hours post injection, which were 5.49 ±2.02% and 5.31±1.20% positive for LNPs, respectively (Figure 2E). These results demonstrate that the intrahepatically injected LNPs were not immediately taken up by the liver, and quickly entered the blood circulation and reached all parts of the body through circulation. The percentage of Cy7 positive cells was highest in the heart and liver, presumably due to their rich blood supply during the fetal period.

The biodistribution of LNPs injected into the developing fetus was significantly different from adult animals. The Cy7-labeled LNPs injected during adulthood were exclusively taken up by the liver, 24 hours post injection. This is consistent with numerous studies demonstrating that LNPs injected during adulthood are mainly captured by macrophages and hepatocytes in the liver [32] and have low efficiency of extrahepatic uptake. In contrast, the unique developing fetal organs during the gestational period [33] allows LNPs delivered at the *in utero* stage to persist in the blood circulation for an extended period and transfect internal organs other than the liver. These results highlight that the fetal period may serve as a unique window for treatment of diseases involving non-liver organs.

### LNPs can deliver CRE mRNA and transfect a wide variety of tissues *in utero* after an intrahepatic injection

The biodistribution studies performed above suggest that *in utero* administration of LNPs has the potential to deliver macromolecules to organs outside of the liver. We therefore performed experiments to determine if intrahepatic injection of mRNA/LNP formulations could transfect mRNA *in utero*, and also identified what cell types and organs were transfected and edited. We used Ai9 CRE reporter mice for these experiments to test the CRE mRNA delivery efficiency. This combination was chosen because it generates a permanent td-Tomato expression in cells that received the mRNA, which allows for identification of the cell types that were transfected. Four different cationic lipid formulations and lipofectamine were screened (see Table 1 for details of formulations) and were injected *in utero* on gestational age of E15.5 and then analyzed 48 hours later via whole fetus and organ imaging. The four LNP based formulations contained various ratios of ionizable lipid D-Lin, DOPE, cholesterol and DMG-PEG2000, and were selected because of their ability to efficiently deliver mRNA in adult mice [26].

Figure 3B demonstrates that LNPs containing D-Lin were remarkably effective at transfecting mRNA *in utero* after an intrahepatic injection, compared to commercially available lipofectamine 2000 which was not effective at transfecting fetal organs and was not investigated further. All of the four kinds of LNP formulations containing D-Lin were able to transfect several organs after an intrahepatic injection. In particular, the heart, lung, liver, kidneys, skin and GI tract were td-Tomato positive after injection with LNPs containing D-Lin (Figure 3B and 3C). The broad tissue distribution of CRE mRNA delivery after an intrahepatic injection suggests that the fetal environment is permeable to nanoparticles, where the nanoparticles are able to enter a variety of vasculature and then transfect a variety of tissues. Intrahepatic injection *in utero* gave a much broader tissue distribution and transfection of CRE mRNA, than an intravenous injection in adult mice. The *in utero* transfection levels *in utero* were similar between the heart, lung, liver, kidneys, GI tract, presumably due to diffusion into the vasculature of multiple organs simultaneously, in contrast, intravenous injection of LNPs in adults generates predominantly transfection in the liver [34]. The ability to transfect organs such as the heart, and kidneys is significant as these organs are the targets for a variety of genetic diseases and have been challenging to transfect with either mRNA or edit via non-viral gene editing, due to the low permeability of their vasculature. Intrahepatic delivery of mRNA/LNP complexes has the potential to create a variety of new therapeutic strategies for genetic diseases.

**Figure 3.**
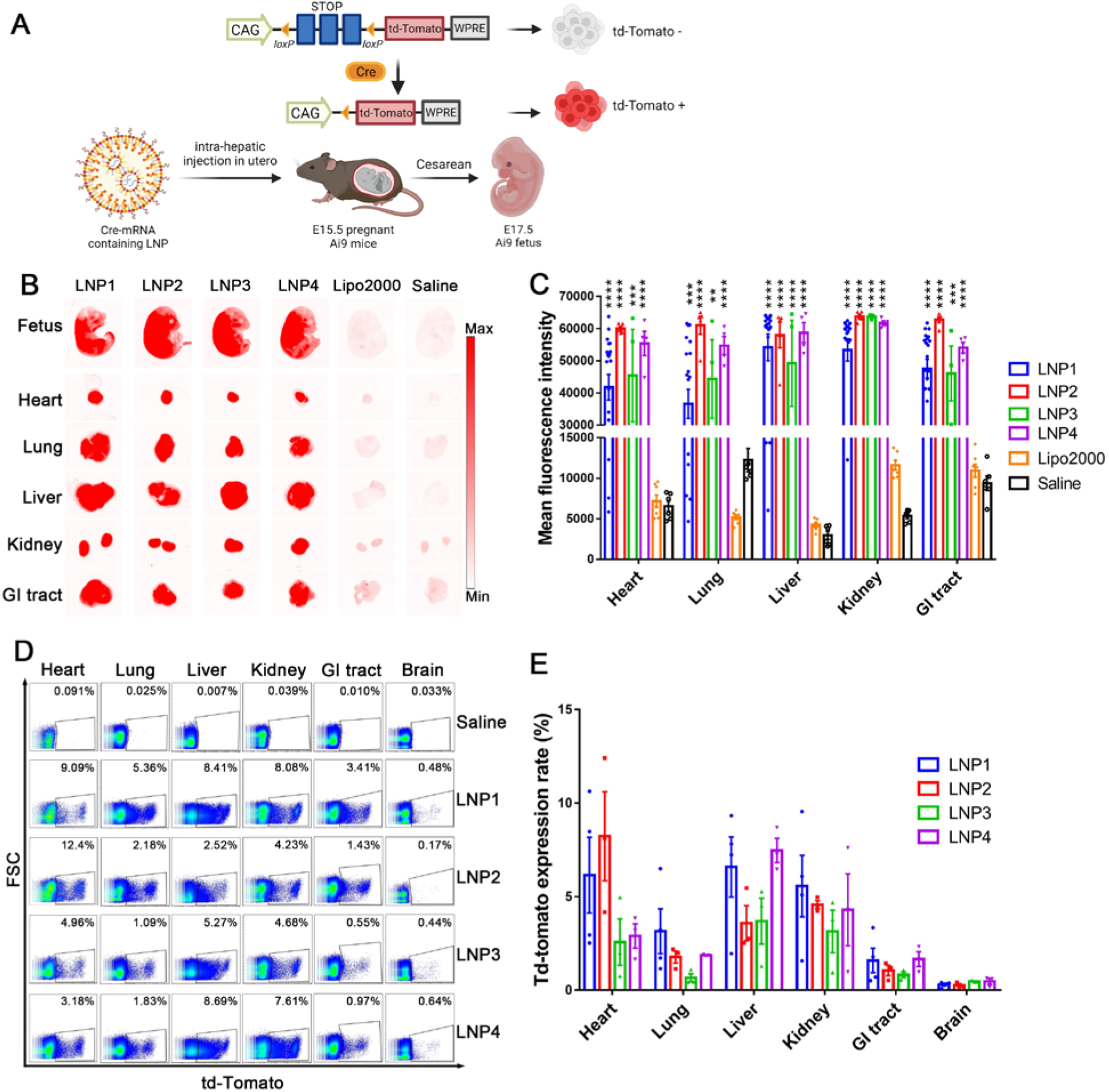
LNPs containing CRE mRNA can transfect the heart, lung, kidney, liver and GI tract *in utero* after an intrahepatic injection. (A) Schematic describing transfection of Ai9 mouse fetuses with CRE mRNA delivered by LNPs 1-4. (B) Whole fetus and internal organ imaging of the transfected mice demonstrate that LNP 1-4 containing D-Lin can efficiently transfect multiple organs at 48 hours post-injection. In particular, the lung, liver, kidney, heart and GI tract were all efficiently transfected. Lipofectamine was not effective after *in utero* injection. (C) Quantitative analysis of the fluorescence observed in B. LNPs were much more efficient at mRNA delivery than lipofectamine (Data are represented as mean ± SEM, n=20 for LNP1, n=5 for LNP2, n=4 for LNP3, n=4 for LNP4, n=7 for Lipo2000). Saline served as a control (n=6). All four LNP formulations edited multiple organs. However, the formulation that contains the highest amount of ionizable lipid (D-Lin) was the most efficient. (D) Flow cytometry analysis of *in utero* transfected organs. (E) Quantitative analysis of D. Greater than 1% editing was observed in a variety of organs. The liver was edited with an efficiency of 8% and the heart and kidney were about 4% (Data are represented as mean ± SEM, n=3).

### LNPs are very efficient at delivering mRNA after an intrahepatic injection *in utero*

The percentage of cells transfected after intrahepatic delivery of mRNA/LNP complexes is a key parameter that will determine the clinical potential of this delivery strategy. Therapeutic gene editing will likely require mRNA delivery efficiencies that are higher than 1%. We consequently determined the mRNA delivery efficiency of mRNA/LNP complexes, containing CRE mRNA. Ai9 mice were treated as described above, and after 48 hours the organs were harvested, digested and analyzed by flow cytometry to determine the percentage of cells that were transfected in each organ. Figures 3D and 3E demonstrate that an intrahepatic injection of mRNA/LNP complexes is remarkably efficient at delivering CRE mRNA. For example, more than 1% of the cells in the heart, lung, liver, kidneys, and GI tract were edited via CRE expression using the intrahepatic delivery route, which appears to have a higher efficiency than the *in utero* intravascular route of administration. To illustrate, Riley et al demonstrated that *in utero* injection of mRNA/LNP complexes via the vitelline vein resulted in 1% of cells in the liver being transfected with GFP [9]. In contrast, intrahepatic injection of LNPs edited approximately 10% of the cells in the liver and 3-4% of the cells in the heart, lung, kidneys, and GI tract in the Ai9 mouse model (Figure 3D and 3E). The blood-brain barrier has probably partially developed and has certain effects even in the fetal period which may explain why the transfection rate of the brain was lower than other organs. It is important to note that the vast majority of these organs have been impossible to edit in adult animals, and *in utero* editing appears to provide a robust methodology for accomplishing this [7, 8, 11].

We further investigated the cell phenotype of transfected cells after in utero intrahepatic injection by immunohistochemistry staining. The phenotype of td-Tomato expressing cells in the heart (Figures 4-c,f,i), diaphragm (Figures 4-d,g,j) and skeletal muscle (Figures 4-e,h,k) was characterized by IHC, staining for Pax7 a primitive myogenic marker [35], and myogenin a transcriptional regulatory protein involved in skeletal muscle differentiation [36]. Large number of td-Tomato positive cells were found in the heart, diaphragm and skeletal muscle. In addition, staining with the proliferation marker cyclin A, demonstrated a wide distribution of proliferating cells that were also TdTomato positive (Figure 4-i,j,k), which represents muscle stem cells that are actively dividing and will proliferate and differentiate into muscle fibers as the fetus develops.

**Figure 4.**
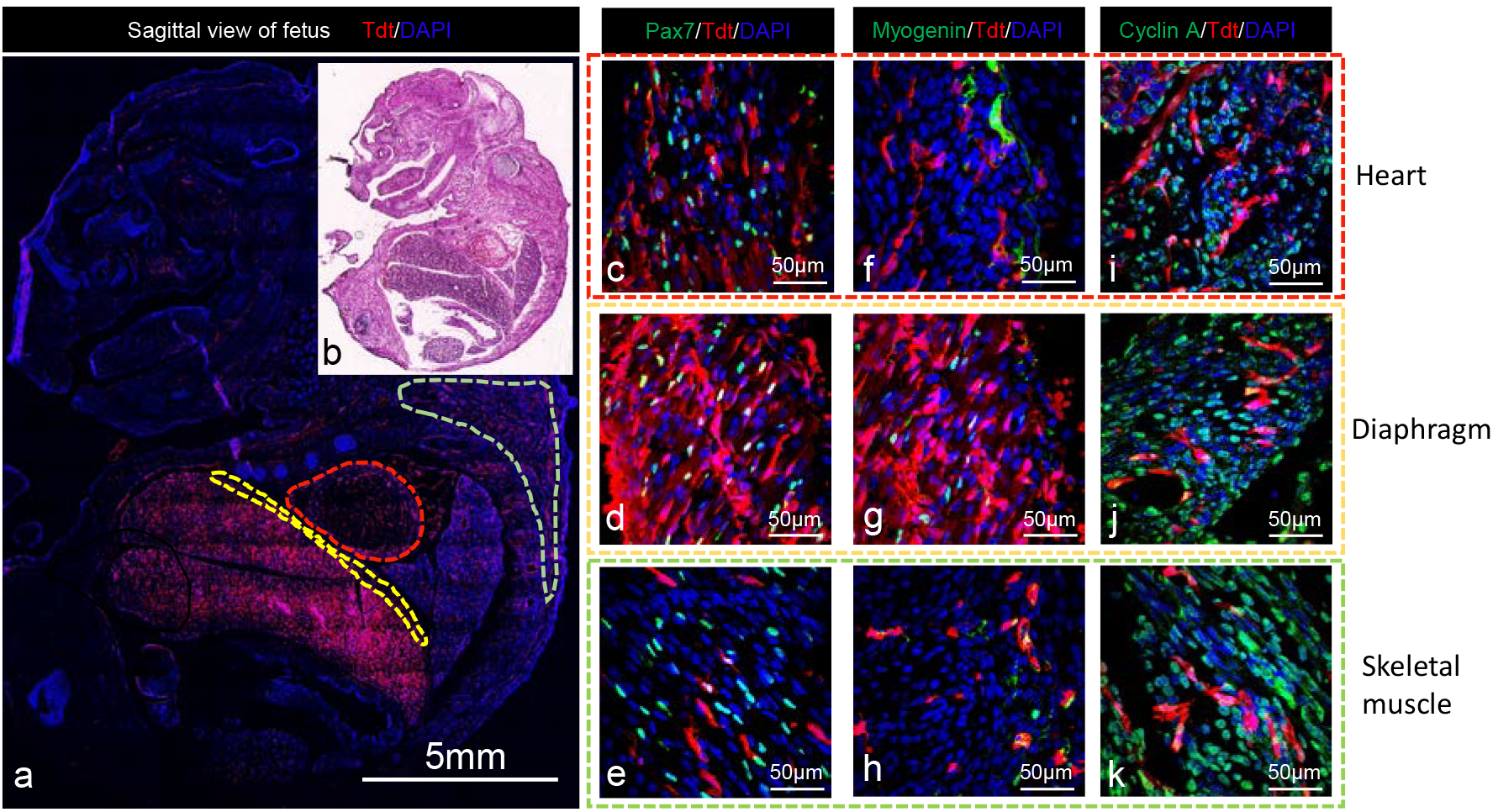
CRE mRNA/LNPs can transfect large number of muscle stem cells in the heart, diaphragm and skeletal muscle. Fluorescent imaging (a) and H&E staining (b) of td-Tomato expression in fetuses at 48 hours after in utero injection of CRE mRNA/LNP1. Large numbers of td-Tomato expressing cells were detected in the heart, diaphragm and skeletal muscle. The phenotype of td-Tomato expressing cells in the heart (c,f,i), diaphragm (d,g,j) and skeletal muscle (e,h,k) were characterized by IHC. Transfected cells were positive for the muscle stem cell markers Pax7 (c,d,e) and myogenin (f,g,h) 48 hours after injection. Transfected cells were also positive for the proliferation marker cyclin A (i,j,k).

### *In utero* injection of CRE mRNA/LNP-1 results in no toxicity to internal organs and has minimal effects on inflammatory cytokine expression levels

We next determined the toxicity of CRE mRNA/LNP complexes to fetuses 48 hours post injection. The fetuses showed good tolerance to *in utero* injection of LNPs. The survival rate of fetuses treated with mRNA/LNP complexes was 96.3% (LNPs 1-4), and close to the survival rate of saline injected fetuses (100%) (see Table S2). The morphology of the internal organs, the heart, lung, liver and kidney were also evaluated by H&E staining. The morphology of the internal organs of mRNA/LNP1 treated fetuses showed a normal structure (Figure 5A upper panel), similar to the saline treated fetuses (Figure S2 upper panel), and was without obvious tissue liquefaction, atrophy, or necrosis. At the same time, a high mRNA delivery efficiency was achieved by this formulation. A large number of td-Tomato positive cells were observed in the internal organs and distributed evenly throughout the organ (Figure 5A middle and lower panel). In addition, there were no td-Tomato positive cells in the saline treated fetuses (Figure S2 middle and lower panel). Finally, we also examined the pregnant mother by histology, via H&E staining and td-Tomato expression, for evidence of toxicity and gene editing, after *in utero* injection with CRE mRNA/LNP1 complexes (see Figures S3 and S4). We did not observe any signs of toxicity in the organs of the mother and 100% of the injected mothers survived (11/11 for CRE mRNA/LNP1), demonstrating that the mother tolerates *in utero* injections with CRE mRNA/LNP1 complexes. We also did not observe any td-Tomato positive cells in the mother indicating that the mother was not transfected by this intervention.

We also measured the cytokine levels in the fetal liver after *in utero* injection with CRE mRNA/LNP1 to determine the inflammatory response triggered by LNP1 (see Figure 5B). Figure 5B demonstrates that CRE mRNA/LNP1 injections had a minimal impact on cytokine expression levels. In particular, 9/10 of the cytokines examined were statistically indistinguishable from the saline treated fetuses, including IFN-γ, IL-1β, IL-2, IL-4, IL-5, IL-6, IL-10, IL-12p70, TNF-α. The level of KC/GRO was increased in LNP treated fetuses compared to saline control mice.

**Figure 5.**
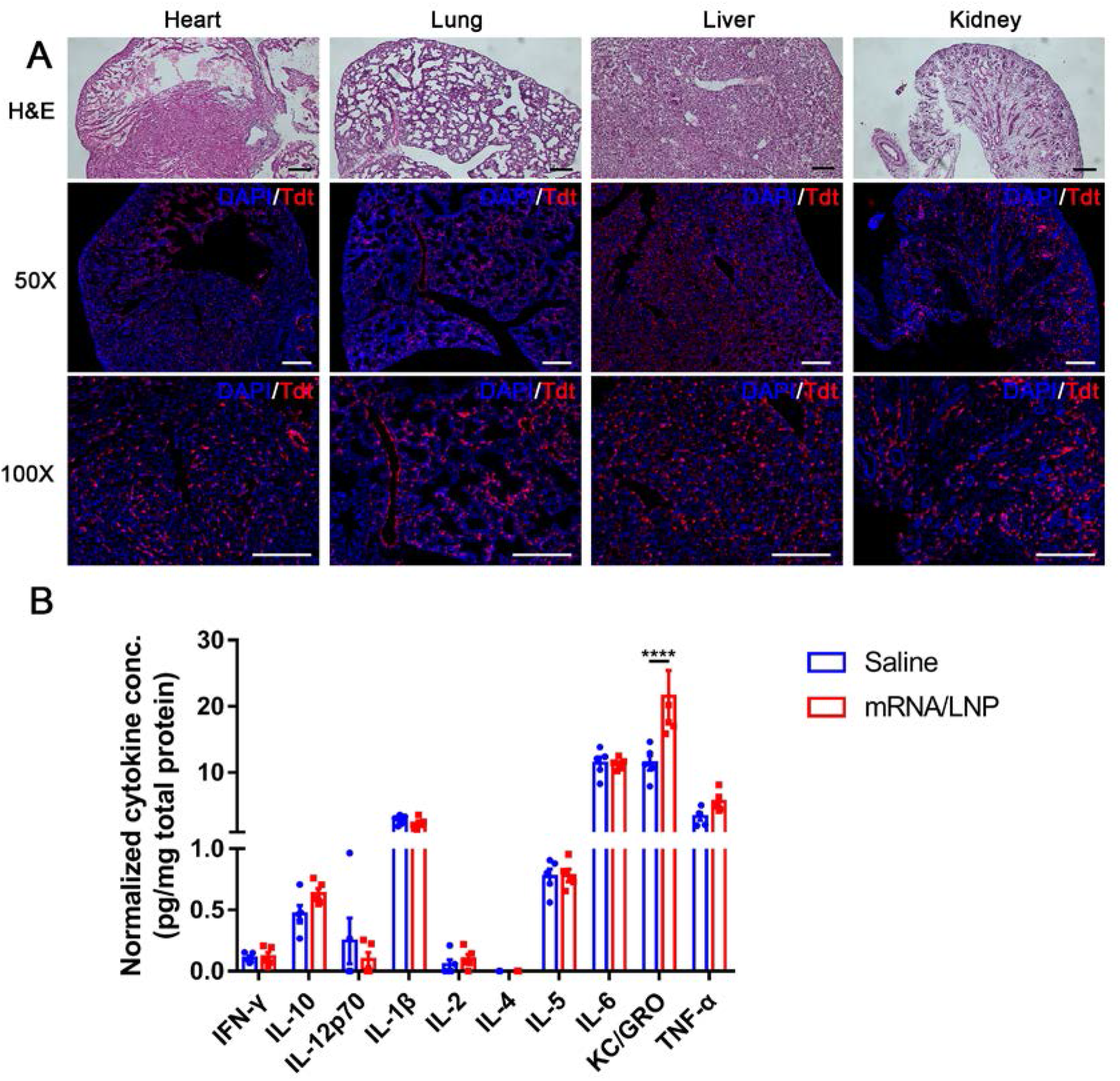
*In utero* injection of CRE mRNA/LNP1 results in no toxicity to internal organs and has minimal effects on inflammatory cytokine expression. (A) H&E staining and fluorescent imaging of td-Tomato expression in fetuses after *in utero* injection of CRE mRNA/LNP1. No signs of tissue damage or toxicity were observed via H&E staining in mRNA/LNP1 treated fetuses. td-Tomato expression was observed throughout the organs and was well distributed (Scale bar = 200 μm). (B) Cytokine analysis of the fetal liver, after *in utero* injection of mRNA/LNP1. Nine of the cytokines analyzed were statistically indistinguishable from the control saline injected mice. KC/GRO had a modest increase in expression (Data are represented as mean ± SEM, n=5).

### LNPs can transfect various cell types in the liver, heart, lungs, kidneys, the GI tract and the brain with high efficiency

A key benefit of using the Ai9 mouse model is that cells transfected with CRE will permanently express td-Tomato and can consequently be identified via flow cytometry. Identifying the cell types transfected with mRNA/LNP formulations will provide valuable information with regards to the type of therapies that can be developed with gene editing therapeutics. LNP1 containing CRE mRNA was injected into Ai9 mouse fetuses as described above. The liver tissue was analyzed for gene editing at different time points after injection. Forty eight hours after injection, the mice were sacrificed, and the internal organ tissues were enzyme dissociated into single cells and stained with a panel of antibodies to determine the editing rates in specific cell populations. Figure 6A and Figure S5 demonstrate that CD90+ cells were the most efficiently transfected cell type in the developing liver, with 55.8 ± 2.52 % of CD90+ cells being transfected. In addition, other important cell types were also transfected, such as CD45+ and CD71+ cells, which are leukocytes and bone marrow progenitors. The transfection rate reached 12.1 ± 1.85 % and 12.17 ± 1.5 % respectively. 34.5 ± 4.77% of cells with endothelial marker CD31+, and 36.13 ± 2.31 % of cells with the epithelial marker CD324+ cells were transfected, which would be sufficient to develop treatments against a variety of genetic diseases associated with these cells. The high level of endothelial cell transfection is presumably because the LNPs are being transported via the developing vasculature and these cells have access to the LNPs more readily than other cells. In adult mice, endothelial cells are difficult to transfect, particularly with LNPs [37]. However, the endothelial cells in the developing fetus should have substantial differences in physiology to adult endothelial cells. In particular they should have a higher rate of cell division and metabolism, and this may make them more phagocytic than adult endothelial cells. Endothelial cells are ideal targets for transfection with secreted proteins. The ability to efficiently transfect endothelial cells in the liver will open up numerous therapeutic applications.

**Figure 6.**
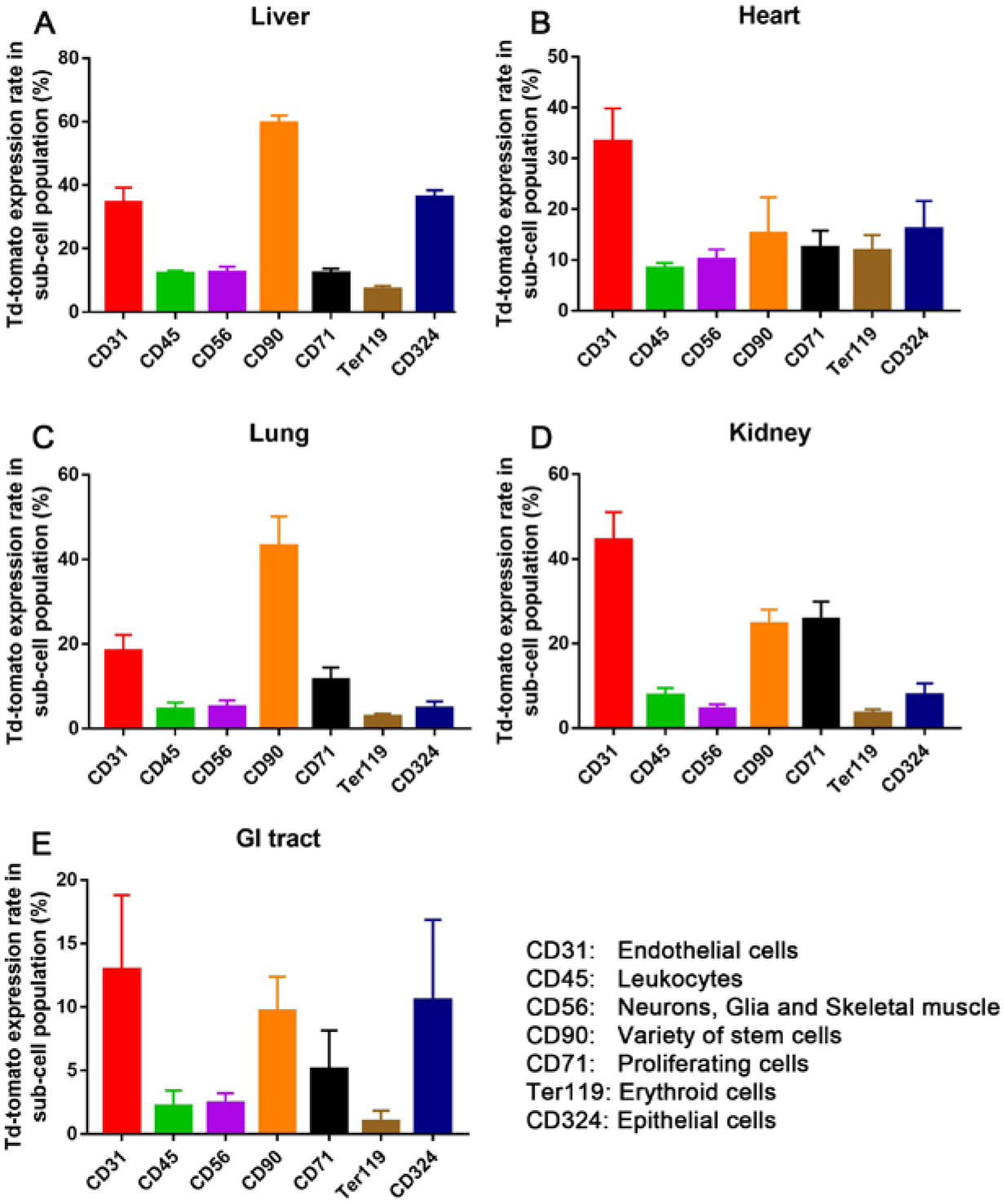
CD31 and CD90 cells are efficiently edited in the fetal heart, lung, kidney and GI tract after an *in utero* injection of mRNA/LNP1 complexes. (A) Flow cytometry analysis of the editing rate in specific cell populations of the liver after in utero injection of CRE mRNA/LNP1 complexes. 60% of CD90 cells were transfected and >30% of CD31 and CD324 cells were also transfected. (B) Cellular analysis of cells transfected in the heart. CD31 cells were transfected with an efficiency of 30% and a variety of other cell types were also transfected with an efficiency > 5%. (C) Cellular analysis of cells transfected in the lung. CD90 cells were transfected with an efficiency of 40% and CD31 were also transfected with an efficiency of 20%. (D) Cellular analysis of cells transfected in the kidney. CD31 cells were transfected with an efficiency of 40%, CD90 and CD71 cells were also transfected with an efficiency of 20%. (E) Cellular analysis of cells transfected in the GI tract. CD31, CD90 and CD 324 cells were transfected with an efficiency of approximately 10%. (Data are represented as mean ± SEM, n=3).

LNPs were able to deliver mRNA to a variety of organs outside of the liver and determining the cell types transfected in these organs will help identify the types of therapeutics that can be generated with *in utero* gene editing. We studied the surface marker expression on the cells that were successfully transfected by mRNA/LNP complexes in the heart, lungs, kidneys, the GI tract and the brain to investigate the phenotypes of the transfected cells. The quantitative analyses are shown in Figure 6 and representative flow cytometry figures are shown in the supplementary figures (Figures S6–S9). The transfection rate of CD31+ cells was the highest in the heart (33.27 ± 6.59 %). While in the lungs, CD90+ cells had the highest transfection rate at 43.1 ± 7.07%. In addition, CD31+ cells also had a relatively high transfection rate (18.37 ± 3.82 %) in the lungs. Similarly, in the kidney, CD31+ cells were the most efficiently transfected cell type in the kidney (44.4 ± 6.61 %), followed by CD90 + cells (24.5 ± 3.52 %) and CD71+ cells (25.73 ± 4.19 %).

### Ai9 mice treated with CRE mRNA/LNP complexes have high levels of editing in the heart and diaphragm in their progeny 4 weeks after birth

A key question with gene editing and mRNA delivery in utero is understanding what organs and cells will be edited in mice that are born and how this editing will persist as the mouse grows and its cells proliferate. The proliferative potential of cells in utero varies dramatically, and how cells edited in utero will map to mice born after development is unknown. This is a key question that needs to be understood before therapeutic gene editing in utero can be accomplished. We performed a series of longitudinal studies in Ai9 mice that were treated with CRE mRNA *in utero,* to determine what cells in the born mice would be edited, after in utero delivery. In particular we wanted to determine if the cells edited in utero had a high proliferative capacity and had the potential to create mosaic organs with large numbers of edited cells. Figure 7 demonstrates that Ai9 mice treated with CRE mRNA/LNP1 complexes have large numbers of edited cells in the heart and diaphragm at 4 weeks after birth in mice. We further stained the heart and diaphragm with the muscle specific antibodies desmin, myosin and laminin (Figure 7A and 7B) to determine if muscle fibers were being edited. The IHC staining demonstrates that a large number of the edited cells at the 4 weeks stage are desmin+, myosin+ or laminin+ muscle fibers, suggesting that edited MuSCs proliferate and differentiate during development. The muscle fiber editing rates in the heart, diaphragm and skeletal muscle after CRE mRNA/LNP1 in utero treatment was quantified by laminin staining. 50.99 ± 5.05%, 36.62 ± 3.42% and 23.7 ± 3.21% of the muscle fibers in the diaphragm, heart and skeletal muscle were respectively edited (Figure 7C). However, CD31 and α-SMA staining showed that only a small number of edited cells were derived from vascular structures (Figure S10 and S11). This is in contrast to 48 hours in utero analysis of the edited cells, where CD31 positive cells were the dominant cell type. The 4 weeks post birth versus 48 hours cell tropism difference suggests that muscle cells proliferate more robustly than endothelial cells, and this may explain the discrepancy between the 48 hour and 4 week timepoints. In addition, the 4 week analysis also suggests that utero mRNA delivery has great potential for treat devastating diseases such as muscular dystrophy and congenital heart disease, due to its ability to transfect the heart and diaphragm with mRNA efficiently.

**Figure 7.**
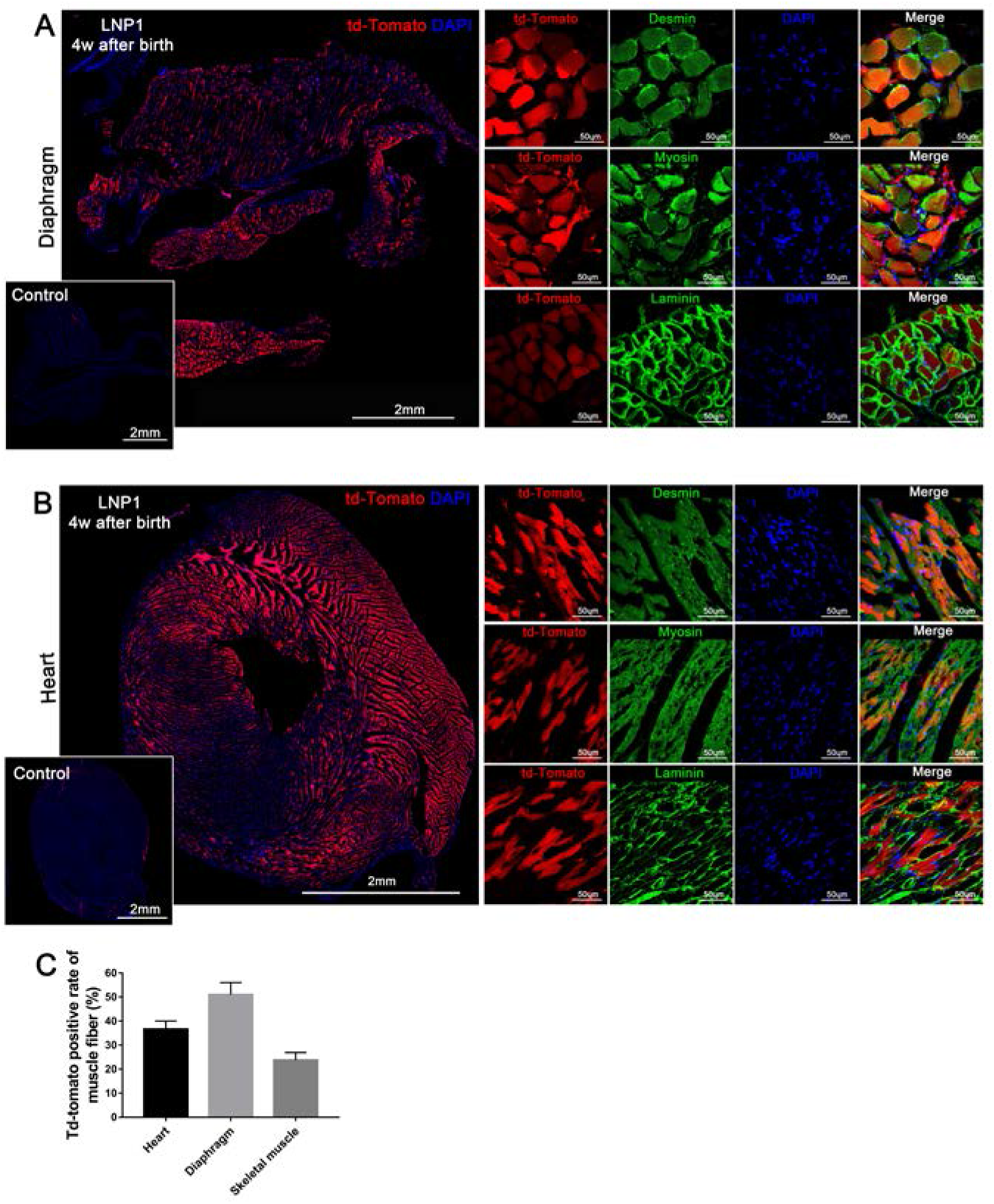
Ai9 mice treated with CRE mRNA/LNP complexes have high levels of editing in the heart and diaphragm in their progeny 4 weeks after birth. Immunohistochemistry staining images of the diaphragm (A) and heart (B). Desmin, Myosin and Laminin staining demonstrates that a large number of mature muscle fibers were edited and expressed td-Tomato 4 weeks after birth. (C) The td-Tomato positive rate of muscle fibers was quantified by Laminin staining. (Data are represented as mean ± SEM, n=4).

### LNPs can transfect Cas9 mRNA and edit internal organs of Ai9 mice

Having demonstrated that an intrahepatic injection of mRNA/LNP complexes was remarkably efficient at delivering CRE mRNA, we next sought to evaluate the efficiency of Cas9 mRNA/sgRNA delivery and its gene editing rate. We performed *in utero* intrahepatic delivery of LNPs containing Cas9 mRNA/sgRNA into Ai9 mouse fetuses following the procedures described above. As described in the Figure 8A experimental scheme, the Cas9 mRNA/sgRNA transduction and genomic excision of the STOP cassette will subsequently activate the expression of td-Tomato. 48 hours after *in utero* injection, the mice were sacrificed and the internal organs including the heart, the liver, lungs, kidneys, the GI tract and the brain were harvested, digested and analyzed by flow cytometry to determine the percentage of cells that were td-Tomato positive at 48 hours post-injection. As shown in Figures 8B and 8C, the Cas9-containing LNPs successfully activated td-Tomato expression in the kidney and liver of Ai9 mice after *in utero* intrahepatic injection. The flow cytometry quantitative analysis in Figure 8C shows that the editing efficiency of LNPs for kidneys and liver is above background levels, and LNP3 activated td-Tomato expression in 0.47 ± 0.14% (n=5) of total kidney cells. These results demonstrate that mRNA/LNP complexes can perform non-viral CRISPR-Cas9 gene editing *in utero.* Compared to the CRE mRNA delivery, the Cas9 gene editing efficiency is relatively low. However, for many enzyme deficiencies therapeutic effects can be achieved even when editing rates are less than 1% [38]. Moreover, the editing efficiency probably will increase during fetal development due to the survival advantage and subsequent expansion of edited cells [39, 40]. Gene editing efficiency is a key problem in in vivo gene therapy, and many new techniques have recently been reported, such as improving the secondary structure of the gRNA or chemically modified the gRNA [41–43], which may also improve the efficiency of in utero gene editing.

**Figure 8.**
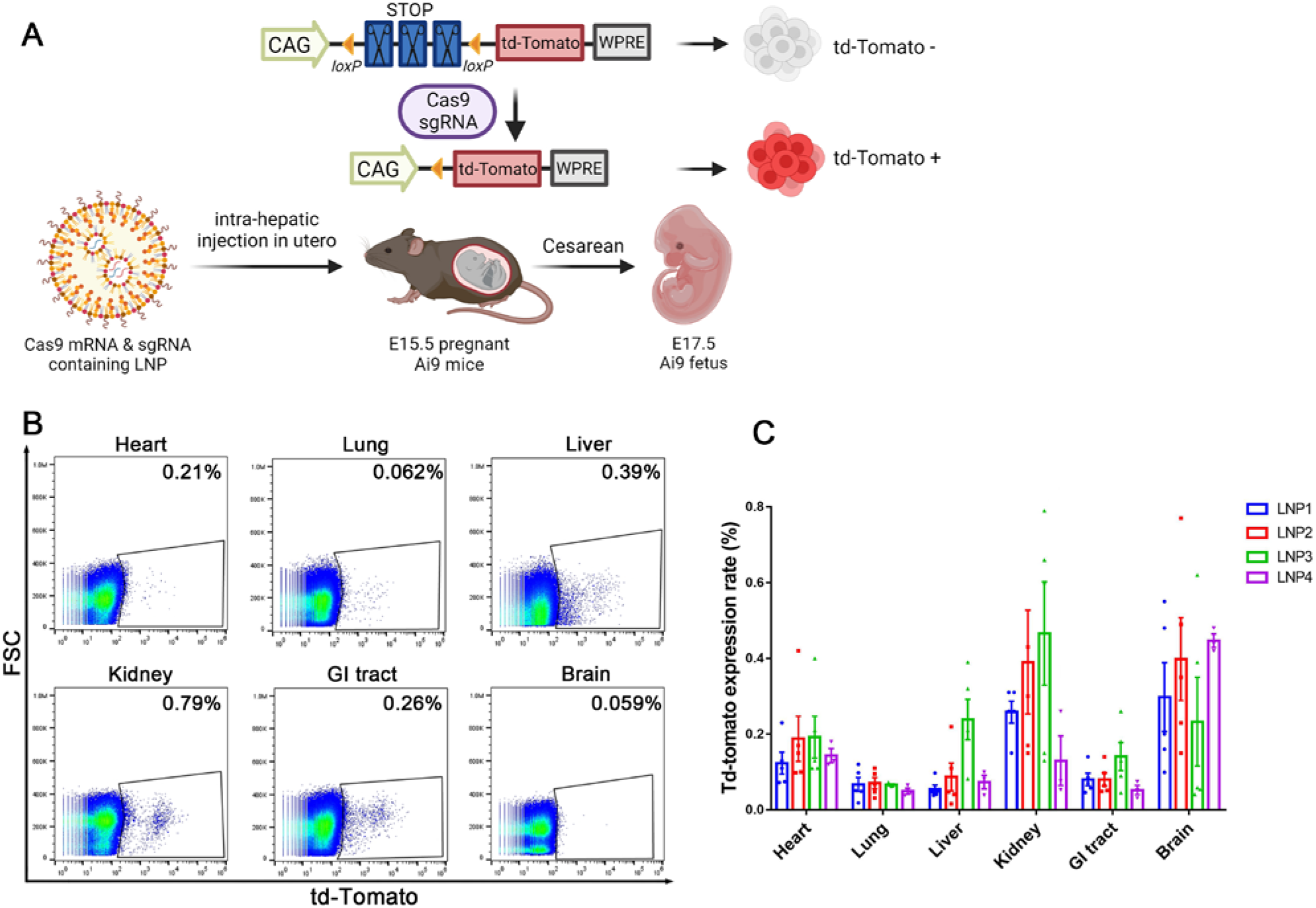
mRNA complexed with LNPs 1-4 containing Cas9 mRNA can edit multiple internal organs after an *in utero* injection. (A) Schematic depicting the procedure of *in utero* Cas9 gene editing in the Ai9 mouse. (B) Flow cytometry of the various organs after transfection with Cas9mRNA/sgRNA/LNP3 complexes. The percent transfection of the various organs was determined. (C) Quantitative analysis of in utero editing with LNPs 1-4 in different organs. LNP3 is the most efficient formulation with regards to Cas9 mRNA delivery *in utero* and edited 0.6% of the cells in the kidney. The liver was also transfected and 0.4% of the cells were transfected (Data are represented as mean ± SEM, n=3).

### Conclusion

In this report we demonstrate that LNPs containing the ionizable lipid D-Lin can deliver mRNA to a wide variety of organs and transfect non-phagocytic cells, such as CD45, CD31, CD90 and CD71 cells. These cell types are almost impossible to transfect with nanoparticles in adult mice because they do not have access to the vasculature. However, the *in utero* environment is characterized by intense levels of angiogenesis and cell division, both of which will increase the permeability of the vasculature and the endocytic activity of non-phagocytic cells. We observed more than 5% transfection levels with D-Lin based LNP formulations, and a wide variety of genetic diseases can potentially be treated with this level of mRNA delivery efficiency. However, the D-Lin LNP formulation was optimized for transfecting adult mice, and it is likely that optimization of LNP formulations for the *in utero* environment will lead to large increases in delivery efficiency, making their translation even more likely. In summary, *in utero* nanoparticle drug delivery has great potential for delivering macromolecules to organs outside of the liver and provides a promising strategy for treating a wide variety of devastating genetic diseases.

## Methods

### Materials

DLin-MC3-DMA was purchased from MedKoo Biosciences. DODAP, DODMA, DOPE, DMG-PEG (MW 2000) (DMG-PEG2000) and DSPE PEG(2000)-N-Cy7 were purchased from Avanti Polar Lipids. Cholesterol was purchased from Sigma Aldrich. Pur-A-Lyzer Midi Dialysis Kits (WMCO, 3.5 kDa) were purchased from Sigma. Cas9 mRNA and Cre mRNA were purchased from TriLink BioTechnologies. Modified sgAi9A, sgAi9B and sgAi9C (Table S3) were purchased from IDT.

### Nanoparticle formation

RNA-loaded LNP formulations were formed using the ethanol dilution method [26]. All lipids with specified molar ratios (Table 1) were dissolved in ethanol and RNA was dissolved in a 10 mM citrate buffer (pH 4.0). The two solutions were rapidly mixed at an aqueous to ethanol ratio of 3/1 by volume (3/1, aq./ethanol, vol./vol.) to satisfy a final weight ratio of 20/1 (total lipids/mRNA), then incubated for 10min at room temperature. All formulations were named based on the additional lipids. The lipids molar ratio for LNP1-3 are D-Lin-MC3-DMA/DOPE/cholesterol/DMG-PEG ratio of 36.8/23.8/38.2/1.2, 45.4/15.7/37.7/1.2, 38.3/11.0/49.5/1.2 and the lipids molar ratio for LNP4 is 35/46.5/16/2.5, follows previous report for D-Lin formulation of mRNA [9]. For biodistribution studies, Cy7 labeled LNPs were used by substitute half of the DMG-PEG2000 in regular LNPs by DSPE PEG(2000)-N-Cy7 in terms of molar ratio. For Cas9/sgRNA, the weight ratio of Cas9 mRNA/sgAi9A/sgAi9B/sgAi9C is 2/0.2/0.2/0.6. After LNP formation, the fresh LNP formulations were diluted with Opti-MEM to 1.25 ng/μl mRNA (with a final ethanol concentration < 0.2 %) for *in vitro* assays and size detection using dynamic light scattering (Zetasizer Nano, Malvern). For RNA encapsulation efficacy was evaluated by the Ribogreen assay[31]. For *in vivo* experiments, the formulations were dialysed (Pur-A-Lyzer Midi Dialysis Kits, WMCO 3.5 kDa) against 1 × PBS for 2 h, and diluted with PBS to 0.125 μg/μl for intrahepatic injections *in utero.*

### Animal studies

All animal procedures were approved by The University of California, Davis (UCD) institutional animal care and use committee (IACUC). All facilities used during the study period were accredited by the Association for the Assessment and Accreditation of Laboratory Animal Care International (AAALAC). Ai9 transgenic mice were purchased from The Jackson Laboratory (Jackson stock No. 007909). Time-mated pregnant Ai9 mice (8-12 weeks old) were bred in-house at UCD and the *in utero* injections were performed as previously described [44]. At E15.5, the mouse was placed under isoflurane anesthesia (5% isoflurane for induction, 2% isoflurane for maintenance) and was positioned supine on a heating pad. A 2-cm midline laparotomy incision was performed and the uterus was carefully exteriorized with cotton applicators. The uterus was examined for the presence of fetuses and the number of normal and nonviable fetuses was noted (the uterus containing fetuses was shown in Figure 2Bi). A glass pipette that had been pulled to a pore size of 70 mm was loaded with 5 μL of CRE mRNA or Cas9 mRNA containing LNPs were resuspended in PBS mixed with blue food coloring at a 1:10 ratio or saline as control. With the aid of a 10X operating microscope, the glass pipette was inserted through the uterus and fetal abdominal wall into the fetal liver. The blue food coloring allowed for visual confirmation of injected reagents located in the liver (Figure 2B ii). The uterus was then returned to its original location in the abdomen and the maternal laparotomy was closed in 2 layers with absorbable 4-0 vicryl suture. The mouse then received a subcutaneous injection of 1 mL PBS for fluid resuscitation and a 0.05 mg/kg dose of buprenorphine for pain management. Mice after *in utero* injection were allowed to carry pregnancy for 48 hours post-treatment and fetuses were delivered by c-section (E17.5). The survival of fetuses was assessed by observation of the size, skin color and spontaneous movement of the fetus. The fetuses and their dissected internal organs including the heart, the liver, lungs, kidneys, the GI tract and brain were imaged by ChemiDocTM MP imaging system (Bio-Rad, Hercules). To quantify the fluorescence density of the internal organs, regions of interest (ROIs) were drawn around the edge of organs and the average fluorescence densities were measured by ImageJ software.

### Biodistribution

The biodistribution experiments were performed on Ai9 mouse fetuses at E15.5 and 12 weeks old adult Ai9 mice. Fetus mice were injected intrahepatically with 5ul of Cy7-labeled LNP1s as described above. 150ul of Cy7 labeled LNP1s were injected into adult mice from inner canthus. The same volume of saline without LNPs were injected as a control. Pregnant mice were euthanized at 3h, 6h and 24h post LNP injection. Fetuses were delivered via cesarean section and washed in PBS. Internal organ including the heart, the liver, lungs, kidneys, the GI tract, and the brain were harvested and imaged by ChemiDocTM MP imaging system (Bio-Rad, Hercules). Adult mice were euthanized at 24h post LNP injection. These mice were perfused with PBS to drain blood. Internal organs were harvested for flow cytometry analyses.

### Flow cytometry

Flow cytometry was used to quantify the percentage of cells in organs that were Cy7 positive or td-Tomato positive following treatment. Single-cell suspensions were obtained as described previously [45, 46] for further staining and flow cytometry analysis. Freshly dissected the heart, the liver, lungs, kidneys, the GI tract, and the brain were minced and digested with 500 μL of 1mg/mL collagenase type I (Gibco) by incubating at 37°C, 5% CO2 for 20 min in 1.5 ml Eppendorf tubes. Cell suspension was collected, neutralized with a medium containing 10% fetal bovine serum (FBS), and placed on ice. Fresh collagenase solution was added to the remaining tissue and incubated for an additional 20 min. Cell suspensions were filtered through 70um Nylon cell strainer (Falcon, cat. 352350) to create a single-cell suspension. The Attune NxT Flow Cytometer (Thermo Fisher Scientific) was used for performing flow cytometry, and FlowJo software (FlowJo LLC) was used for data analyses. All antibodies were obtained from BD Biosciences. Cells were stained with BV421-CD56 (cat.748094), APC-CD31 (551262), FITC-CD90.2 (553003), BV421-CD45 (563890), BV650 CD324 (752472), FITC-CD71 (561936), APC-Ter119 (557909) from BD Biosciences. BDTM anti-Rat Ig, *κ* CompBeads (552844) were used to generate compensation controls.

### Histology and immunofluorescence staining

Fetal and maternal internal organs were fixed in 4% PFA for 24 h, dehydrated with 30% sucrose for 24 h, and embedded in the O.C.T compound (Sakura Finetek USA). Serial sections were made at the thickness of 10 μm using a Cryostat (Leica CM3050S) and collected onto microscope slides (Matsunami Glass). H&E staining was performed to characterize tissue morphology. Tissue sections were extensively washed with PBS, heat antigen retravel were performed within target retrieval solution, pH 9.0 (Dako). The sections were blocked with 5% BSA in PBS at room temperature for 1 h and stained with primary antibody at 4 °C overnight. The dilutions of primary antibodies were 1:100 for Desmin (abcam, ab32362), 1:200 for Myosin (Millipore Sigma, M1570), 1:100 for Laminin (Millipore Sigma, L9393), 1:100 for CD31 (Novus Biologicals, AF3628) and 1:200 for α-SMA (LSBio, LS-B3933), 1:10 for Pax7 (Developmental Studies Hybridoma Band), 1:100 for Cyclin A (Millipore Sigma, ZRB-1590), 1:50 for Myogenin (Santa Cruz Biotechnology, SC-12732). Sections were incubated with their respective secondary antibodies diluted at 1:250 for 30min at room temperature. The slides were stained with 1:5000 dilution of DAPI for 5 min, mounted with Prolong Diamond Antifade Mountant (Invitrogen). Fluorescent imaging of td-Tomato was performed with a Zeiss Observer Z1 microscope.

### Cytokine analysis

Fetal liver lysates were harvested and assessed for cytokine levels at 48 hours post LNP1 injections. The freshly harvested fetal livers were lysed in Cell Lysis Buffer (Cell signaling technology) supplemented with 1% Protease Inhibitor Cocktail (Sigma-Aldrich). Total protein content in each sample was determined with the BCA protein assay kit (Thermo Fisher Scientific) per manufacturer’s instructions. The cytokine levels were assessed using the Proinflammatory panel 1 (mouse) kit for MSD multi-spot assay system according to the manufacturer’s instructions. Saline injection served as the control. Liver cytokine data were normalized to the total liver protein concentration, as determined by the BCA assay.

### Statistic

Statistical analysis was performed by one-way analysis of variance (ANOVA) following Tukey’s multiple comparison tests using PRISM 7 (GraphPad Software Inc., San Diego, CA, USA) for experiments involving the comparison of more than two groups. The flow cytometry characterized biodistribution and cytokine expression levels were analyzed by student’s t test. Statistical differences were considered significant if p < 0.05.

## Supporting Information

Supporting Information is available from the Wiley Online Library or from the author.

## Acknowledgements

This work was in part supported by the National Institutes of Health grants UG3NS115599, R61DA048444-01, R01MH125979 (N.M.), 1R01NS100761 and 1R01NS115860 (A.W.).

## Supporting Information

**Table S1.** Characterization of CRE mRNA LNPs formulations.

| LNP formulations | Size (nm) | PDI | Encapsulation efficiency (%) |
| --- | --- | --- | --- |
| LNP1 | 144.0 ± 0.8 | 0.114 ± 0.014 | 76.82 ± 0.25 |
| LNP2 | 120.5 ± 15.0 | 0.126 ± 0.019 | 76.27 ± 1.29 |
| LNP3 | 132.0 ± 8.5 | 0.108 ± 0.015 | 82.92 ± 1.75 |
| LNP4 | 112.5 ± 2.9 | 0.085 ± 0.016 | 74.65 ± 0.52 |

LNPs had a hydrodynamic diameters ranging from 112.5-144.0 nm and narrow size distribution. The mRNA encapsulation efficiency is good with a range between 74.65%-82.92% by RiboGreen assay. (Data are represented as mean ± SEM, n=3)

**Figure S1.**
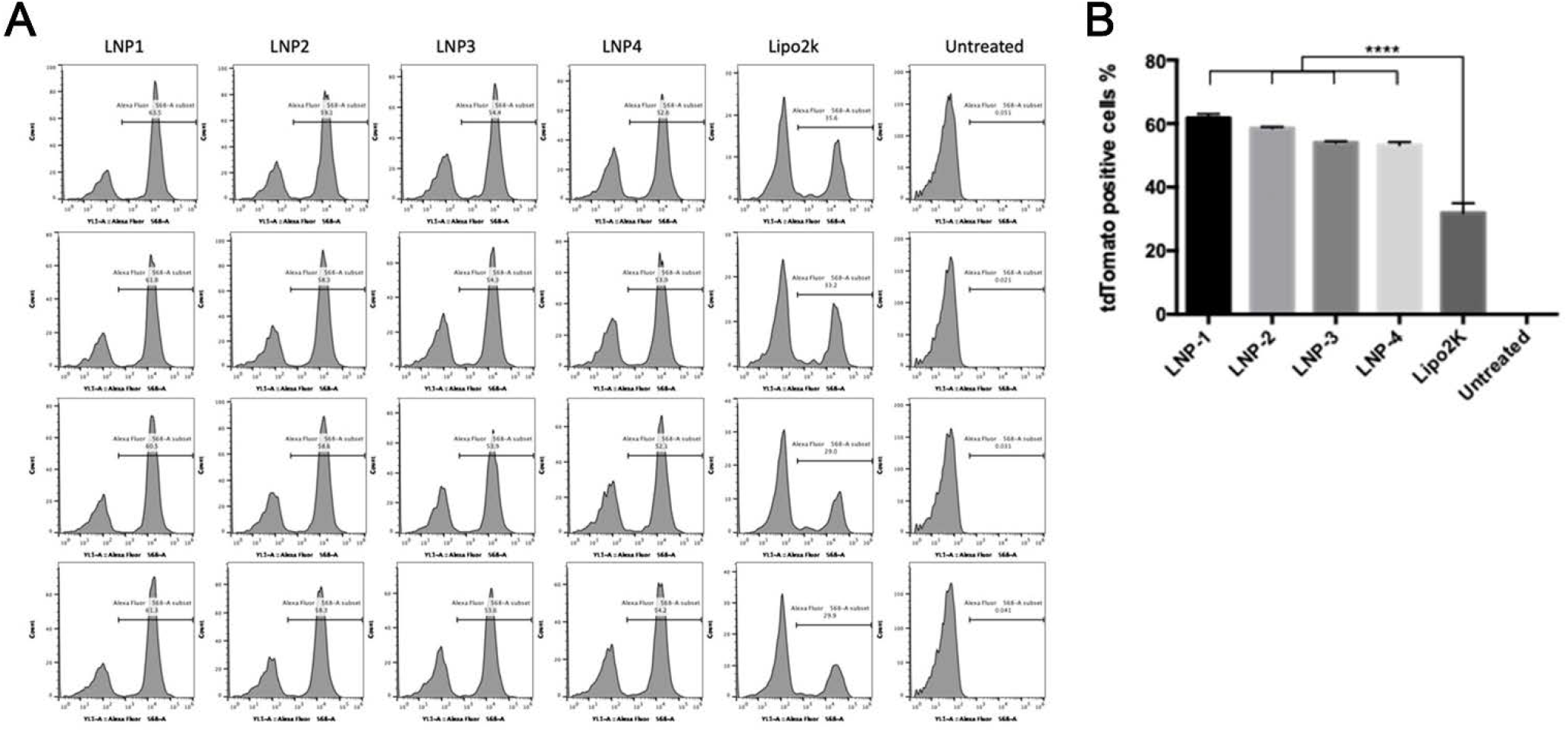
LNPs CRE mRNA formulations showed excellent *in vitro* gene editing efficiency in Ai9 NIH 3T3 cells. LNPs showed significantly higher gene editing efficiency compared with traditional lipofectamine 2000 transfection, with at least 66% increase of td-Tomato positive cells. (Data are represented as mean ± SD, n=4) Detailed flow cytometry data is also showed.

**Table S2.** Survival rate of fetuses 48 hours post in utero injection with CRE mRNA/LNP complexes

|  | N of total injected | N of survival | Survival rate (%) |
| --- | --- | --- | --- |
| LNP1 | 41 | 40 | 97.6 |
| LNP2 | 5 | 5 | 100 |
| LNP3 | 4 | 3 | 75 |
| LNP4 | 4 | 4 | 100 |
| Saline | 9 | 9 | 100 |

**Figure S2.**
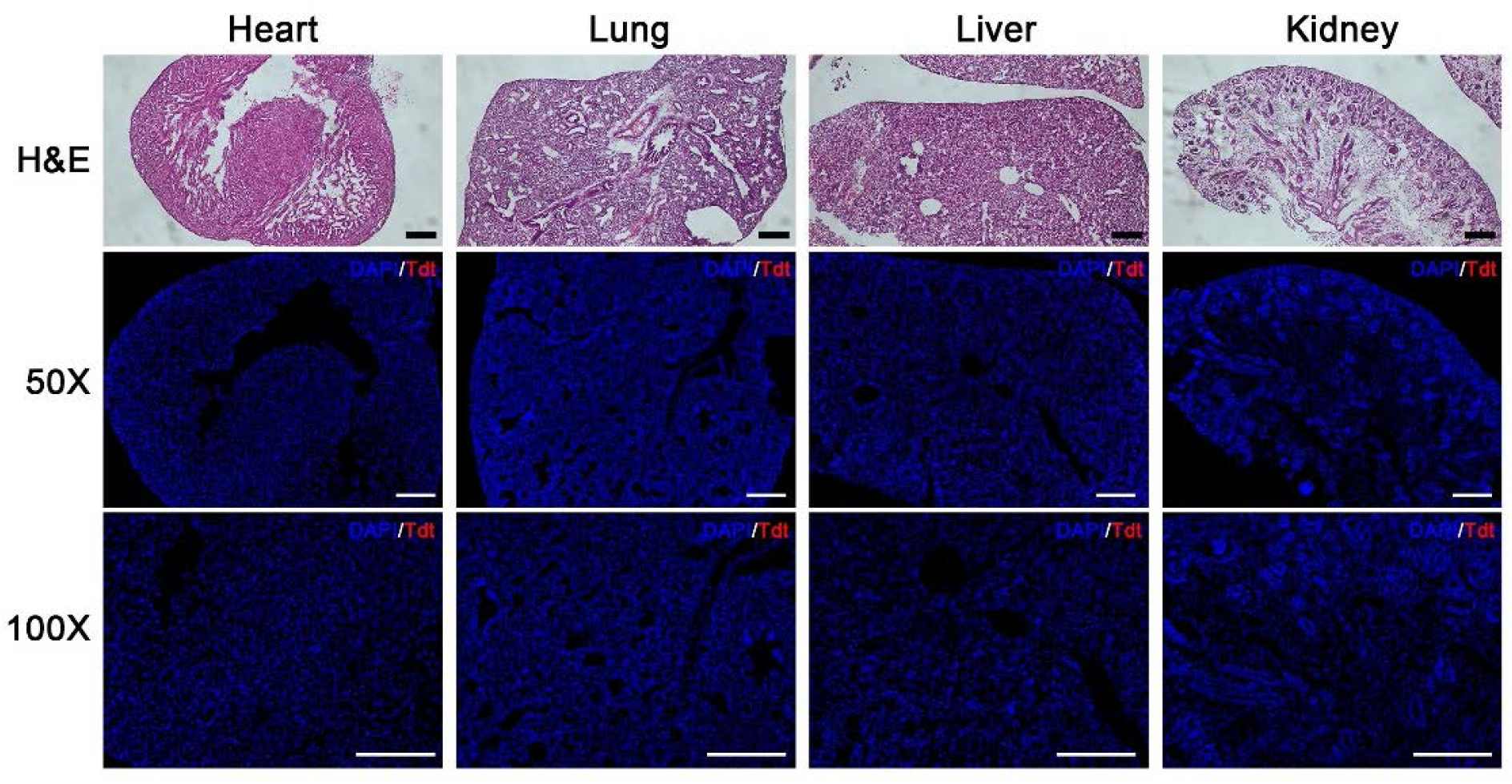
In utero injection of saline results in no toxicity to internal organs and does not cause Td-tomato expression. H&E staining and fluorescent imaging of td-Tomato expression in fetuses after in utero injection of saline. No signs of tissue damage or toxicity were observable via H&E staining in saline treated fetuses. td-Tomato expression was not observed in the organs. (Scale bar = 200 μm)

**Figure S3.**
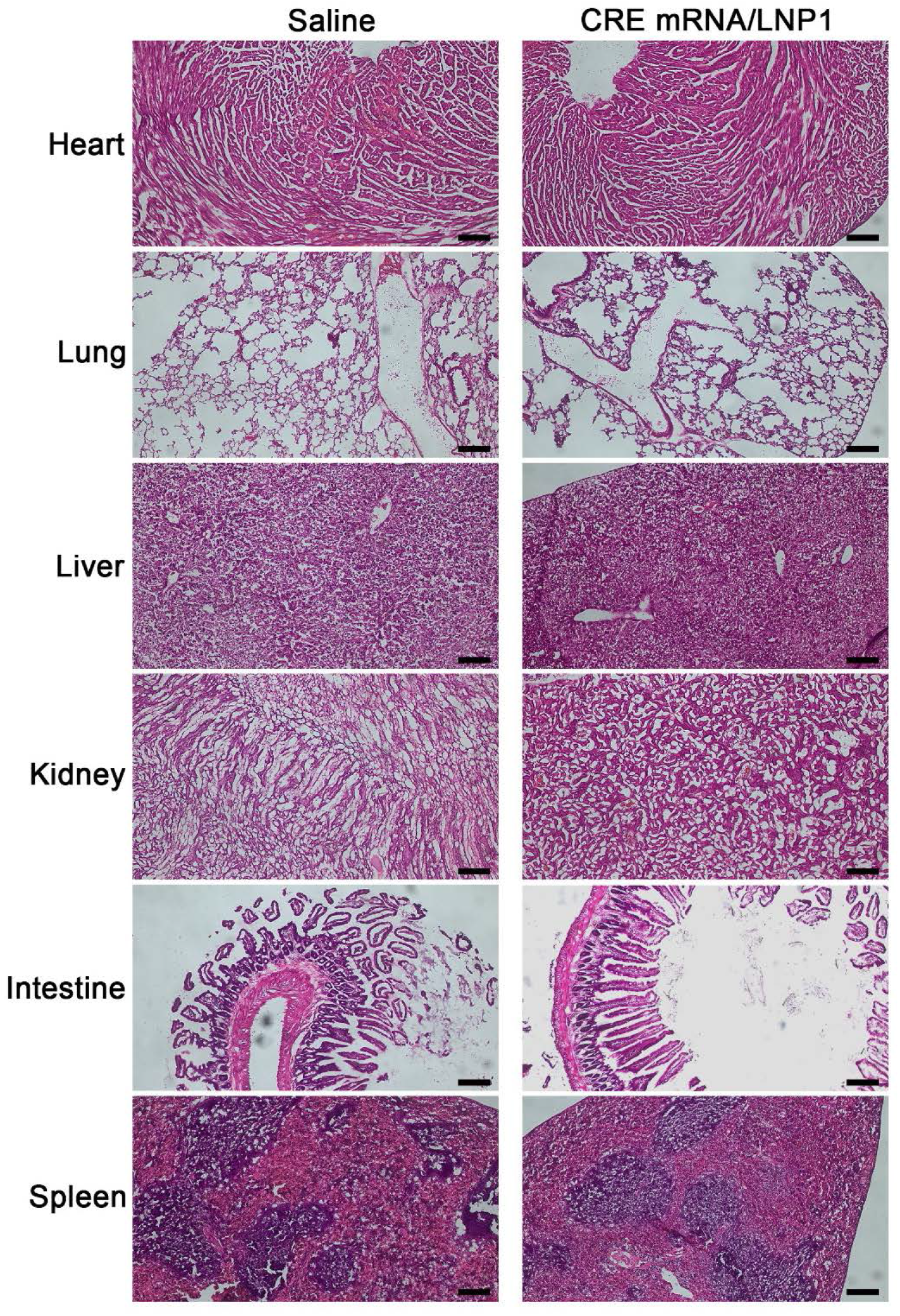
In utero injection of CRE mRNA/LNP1 results in no toxicity to internal organs of the mother. H&E staining of the internal organs of the mother demonstrate that in utero injection of CRE mRNA/LNP1 does not cause tissue damage. (Scale bar = 200 μm)

**Figure S4.**
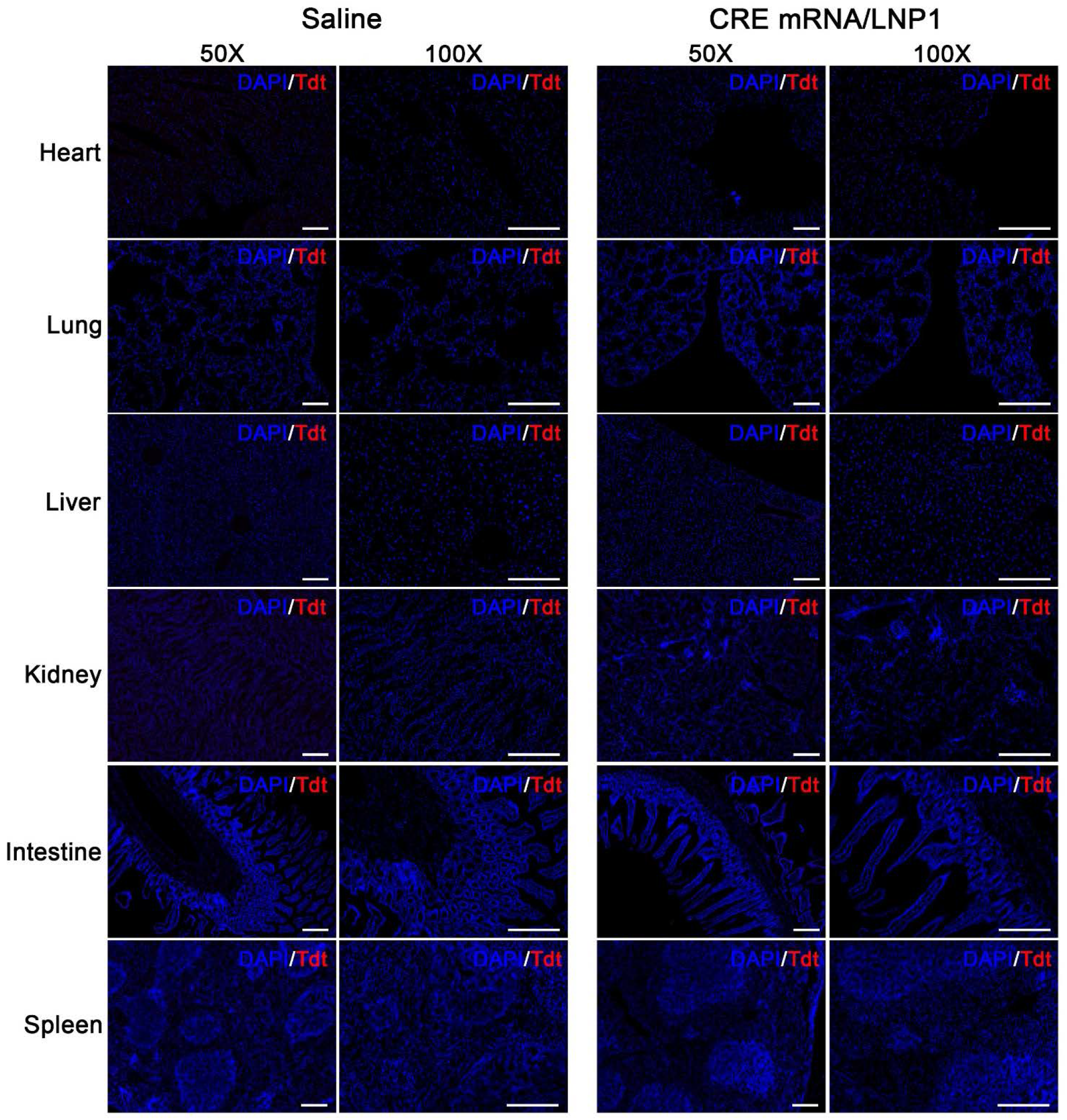
In utero injection of CRE mRNA/LNP-1 does not induce td-Tomato expression in the mother. Fluorescent imaging of Td-tomato and DAPI of the internal organs of the mother demonstrate that in utero injection of CRE mRNA/LNP1 does not induce td-Tomato expression in the mother. (Scale bar = 200 μm)

**Figure S5.**
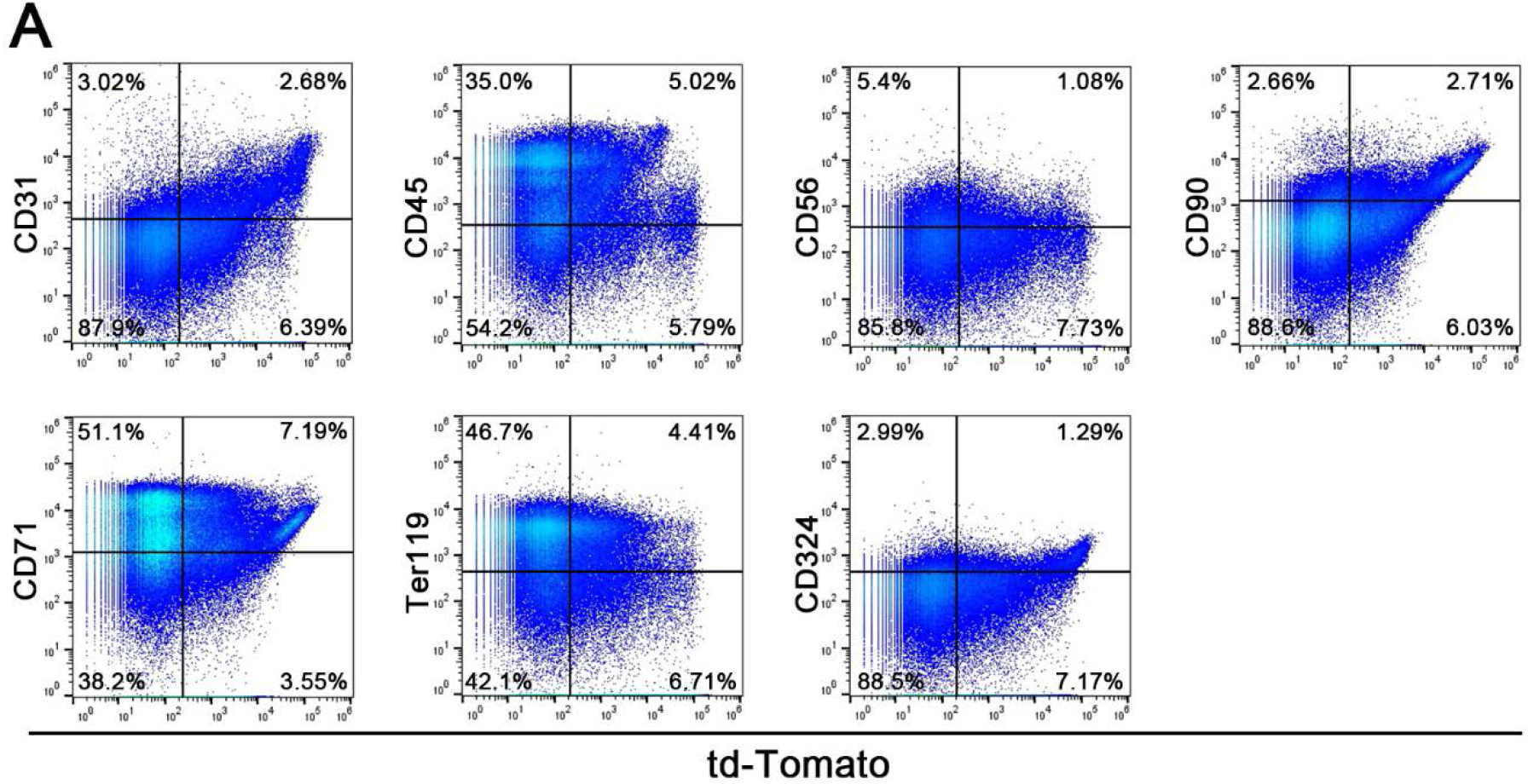
mRNA/LNP1 complexes can edit a variety of cell types in the liver. Representative flow cytometry analysis of the heart cells after transfection with Cre-mRNA/LNP1 complexes. The quantitative analysis was shown in Figure 6A.

**Figure S6.**
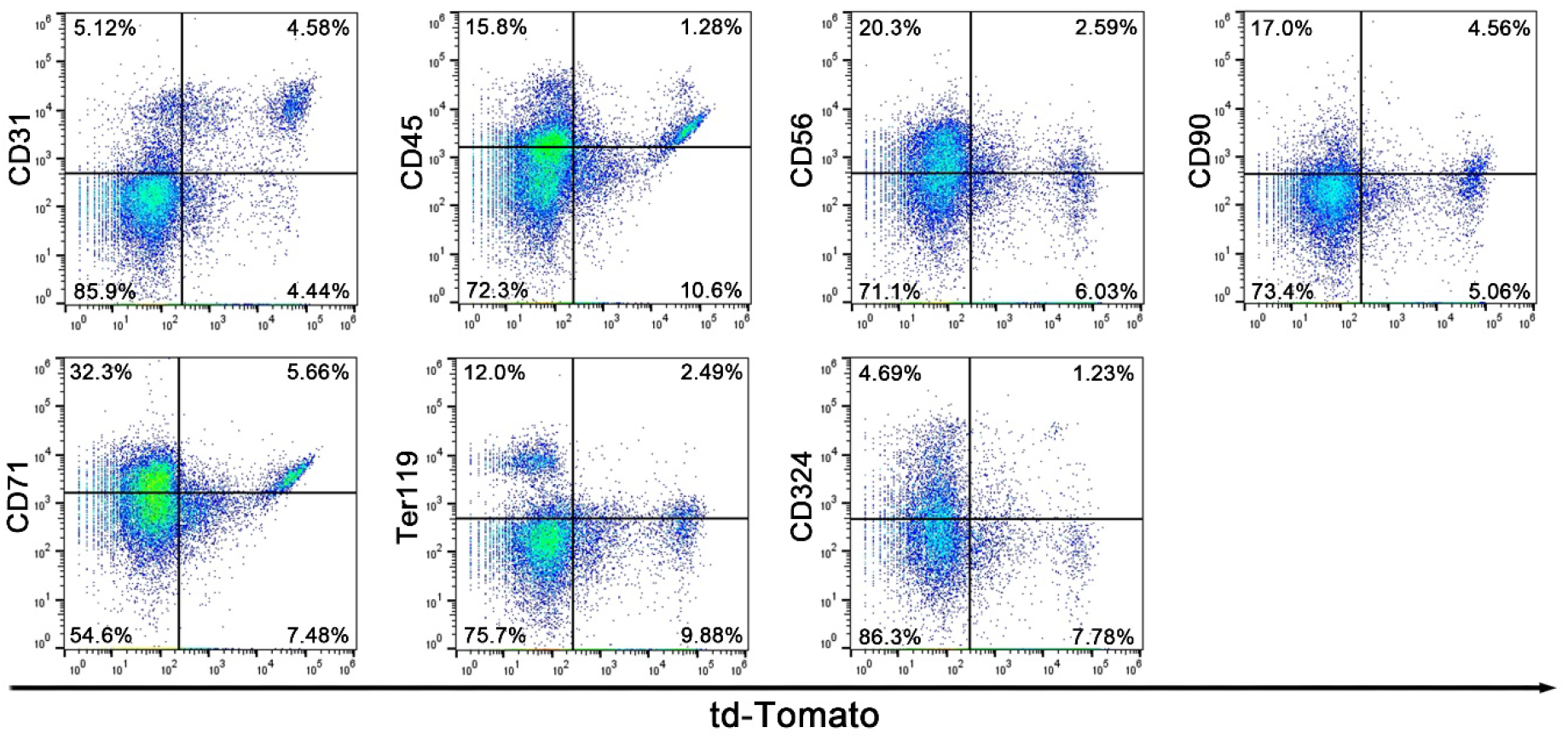
mRNA/LNP1 complexes can edit a variety of cell types in the heart. Representative flow cytometry analysis of the heart cells after transfection with Cre-mRNA/LNP1 complexes. The quantitative analysis was shown in Figure 6B.

**Figure S7.**
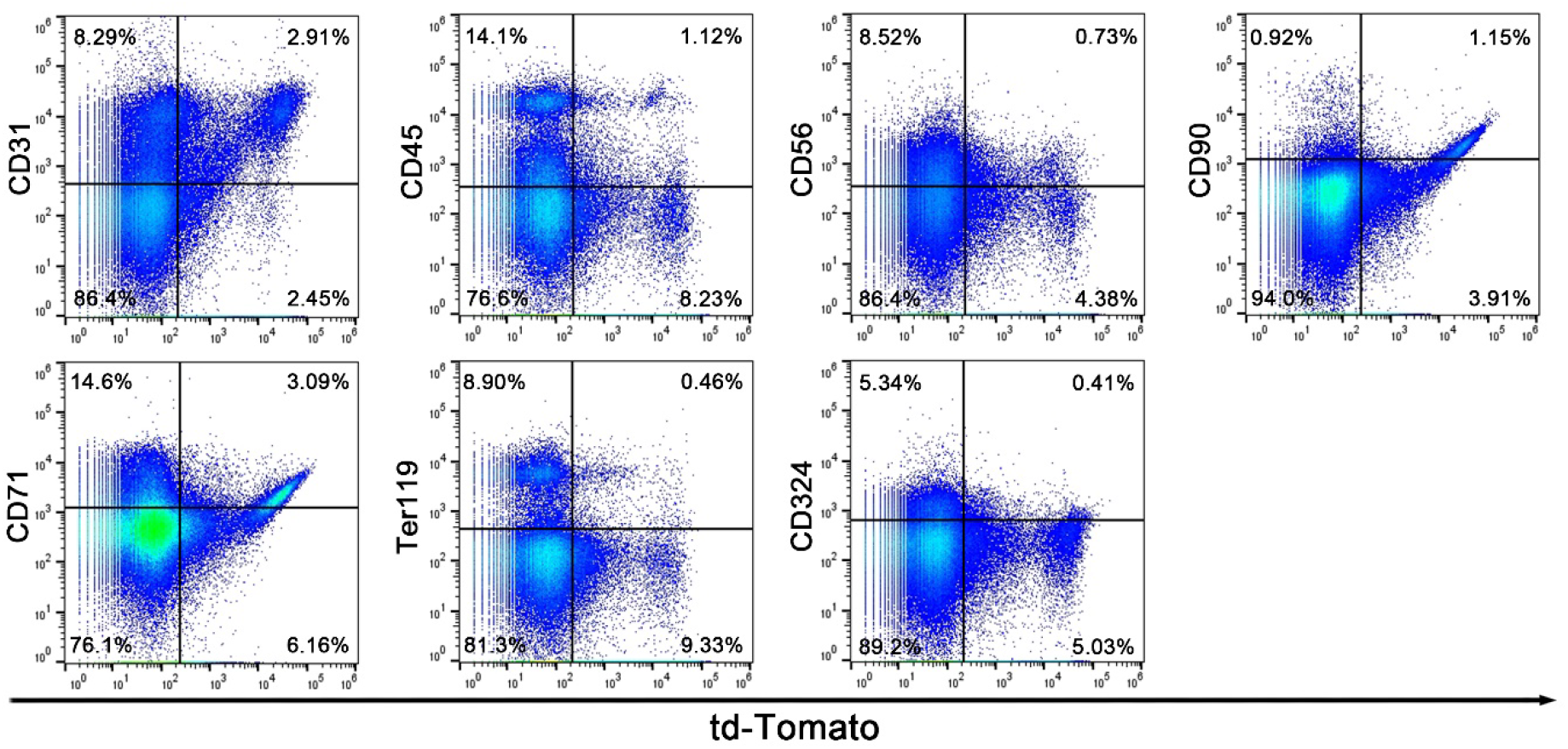
mRNA/LNP1 complexes can edit a variety of cell types in the lung. Representative flow cytometry analysis of the lung cells after transfection with Cre-mRNA/LNP1 complexes. The quantitative analysis was shown in Figure 6C.

**Figure S8.**
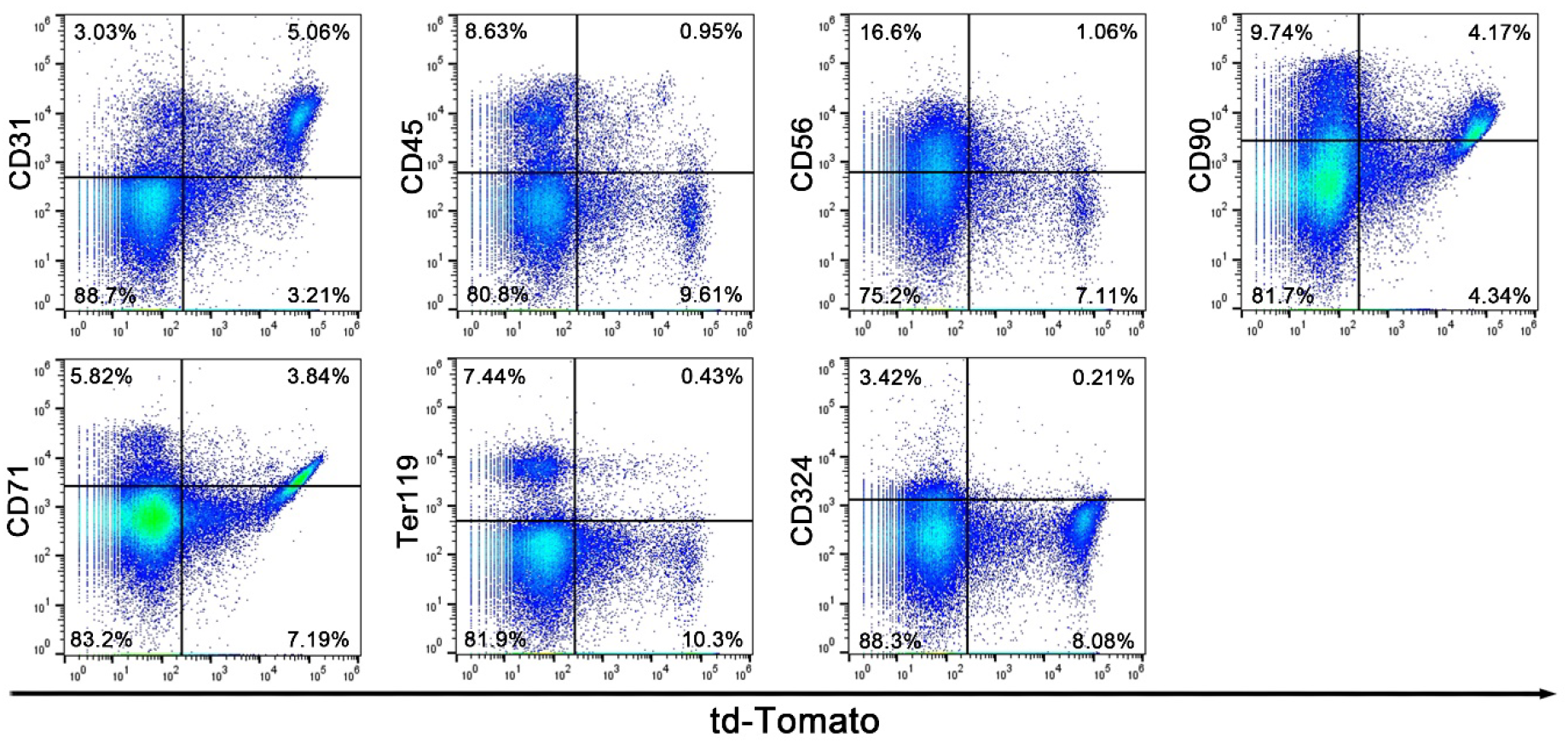
mRNA/LNP1 complexes can edit a variety of cell types in the kidney. Representative flow cytometry analysis of the kidney cells after transfection with Cre-mRNA/LNP1 complexes. The quantitative analysis was shown in Figure 6D.

**Figure S9.**
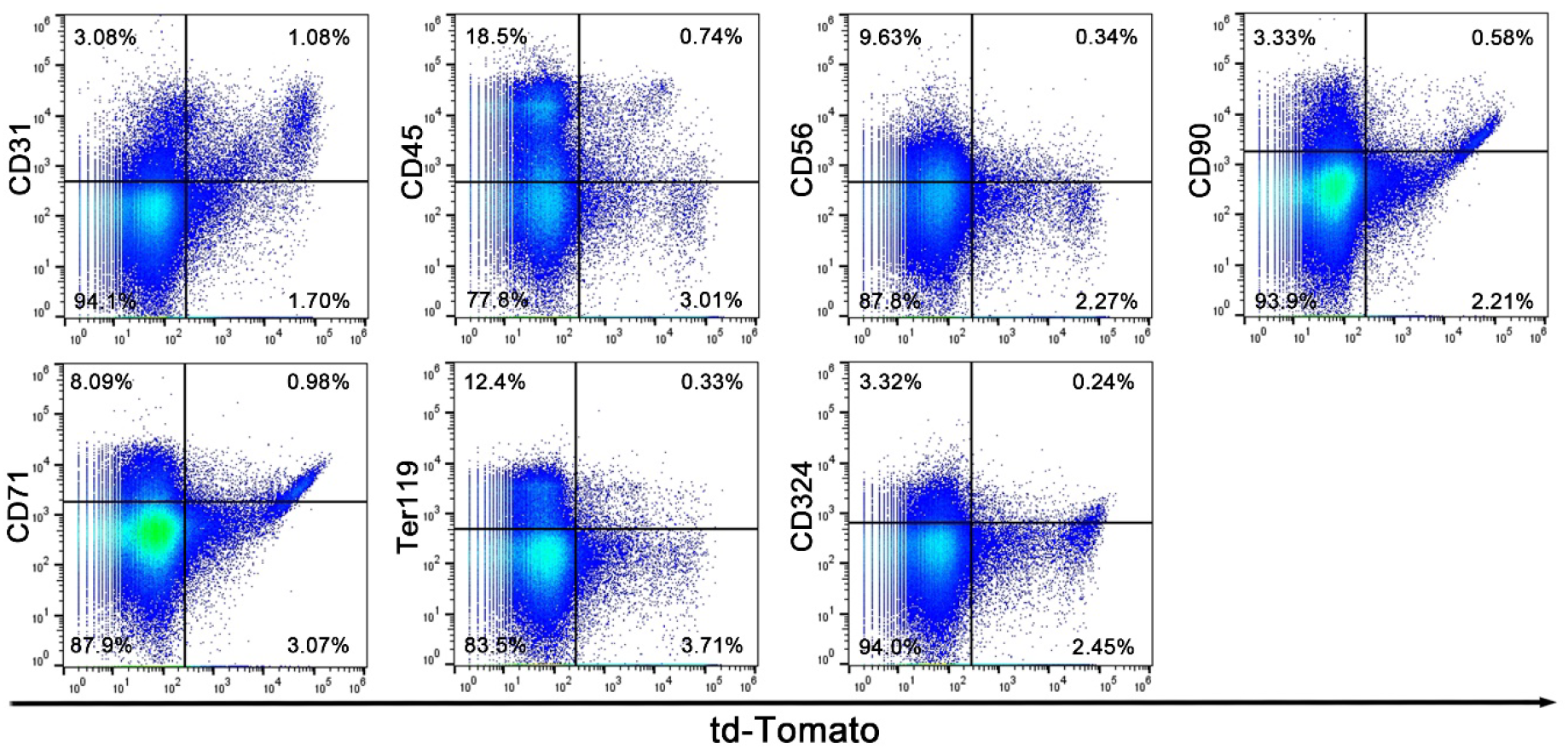
mRNA/LNP1 complexes can edit a variety of cell types in the GI tract. Representative flow cytometry analysis of the cells in GI tract after transfection with Cre-mRNA/LNP1 complexes. The quantitative analysis was shown in Figure 6E.

**Table S3.**
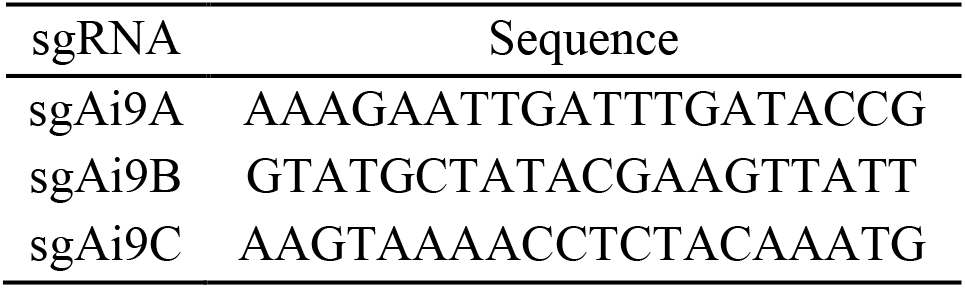
sgRNA protospacer sequence

**Figure S10.**
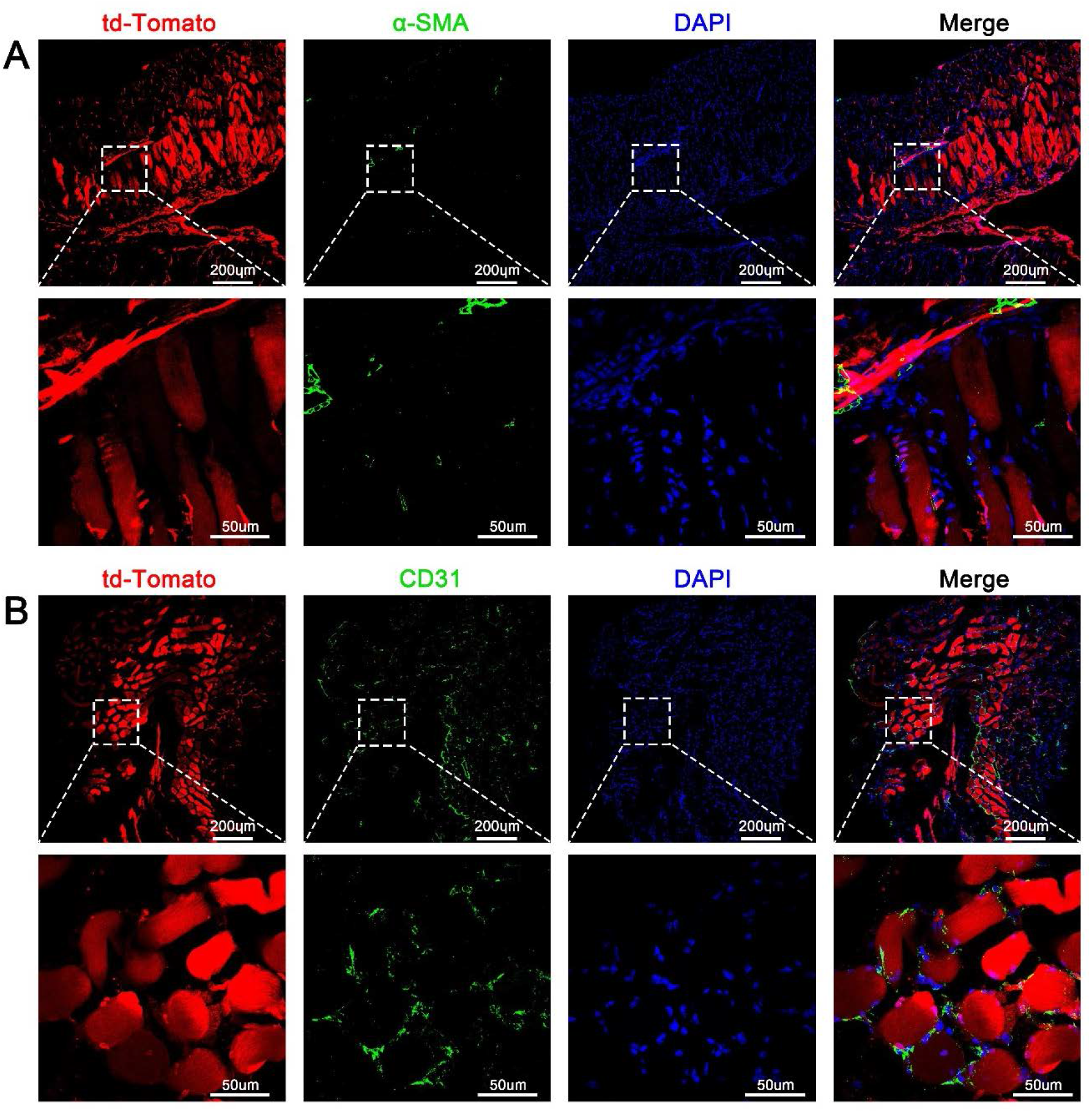
The td-Tomato expression cell rate in internal organ of CRE mRNA/LNP1 in utero injected mice were maintained at 1 day and 4 weeks after birth. Representative flow cytometry analysis of the in utero transfected organs at 48h after injection, 1 day and 4 weeks after birth respectively. (B) Quantitative analysis of A. (Data are represented as mean ± SEM, n=6 for 48h post injection, n=4 for 1day after birth and 4 weeks after birth)

**Figure S11.**
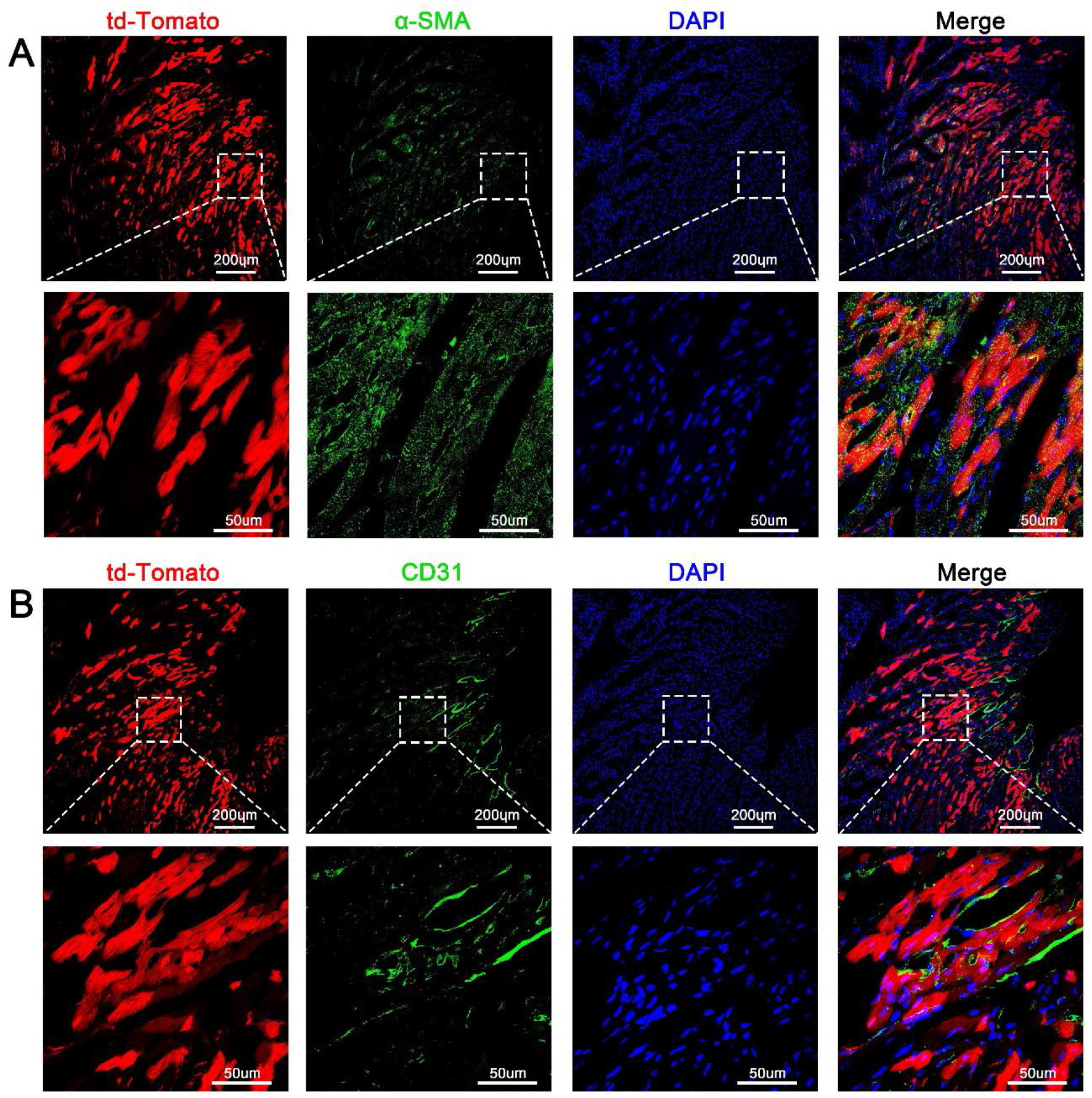
The td-Tomato expression cell rate in internal organ of CRE mRNA/LNP1 in utero injected mice were maintained at 1 day and 4 weeks after birth. (A) Representative flow cytometry analysis of the in utero transfected organs at 48h after injection, 1 day and 4 weeks after birth respectively. (B) Quantitative analysis of A. (Data are represented as mean ± SEM, n=6 for 48h post injection, n=4 for 1day after birth and 4 weeks after birth)

